# Single-Cell Data Integration and Cell Type Annotation through Contrastive Adversarial Open-set Domain Adaptation

**DOI:** 10.1101/2024.10.04.616599

**Authors:** Fatemeh Aminzadeh, Jun Wu, Jingrui He, Morteza Saberi, Fatemeh Vafaee

**Author notes:** Equal first Authorship (Fatemeh Aminzadeh, Jun Wu). Corresponding Author Correspondence to: A/Professor Fatemeh Vafaee, School of Biotechnology and Biomolecular Sciences, UNSW SYDNEY NSW 2052 AUSTRALIA, E.

## Abstract

Single-cell sequencing technologies have enabled in-depth analysis of cellular heterogeneity across tissues and disease contexts. However, as datasets increase in size and complexity, characterizing diverse cellular populations, integrating data across multiple modalities, and correcting batch effects remain challenges. We present SAFAARI (Single-cell Annotation and Fusion with Adversarial Open-Set Domain Adaptation Reliable for Data Integration), a unified deep learning framework designed for cell annotation, batch correction, and multi-omics integration. SAFAARI leverages supervised contrastive learning and adversarial domain adaptation to achieve domain-invariant embeddings and enables label transfer across datasets, addressing challenges posed by batch effects, biological domain shifts, and multi-omics modalities. SAFAARI identifies novel cell types and mitigates class imbalance to enhance the detection of rare cell types. Through comprehensive benchmarking, we evaluated SAFAARI against existing annotation and integration methods across real-world datasets exhibiting batch effects and domain shifts, as well as simulated and multi-omics data. SAFAARI demonstrated scalability and robust performance in cell annotation via label transfer across heterogeneous datasets, detection of unknown cell types, correction of batch effects, and cross-omics data integration while leveraging available annotations for improved integration. SAFAARI’s innovative approach outperformed competing methods in both qualitative and quantitative metrics, offering a flexible, accurate, and scalable solution for single-cell analysis with broad applicability to diverse biological and clinical research questions.

## Introduction

The rapid advancements in single-cell technologies across diverse omics modalities have revolutionized our understanding of cell biology and human disease by allowing in-depth analysis of cellular heterogeneity across tissues, tumors, organs, and entire organisms. However, as single-cell datasets continue to expand in both size and complexity, new computational challenges arise, particularly in managing increased scale, extended modalities, and inevitable batch effects. Deep learning-based approaches have proven instrumental in modeling relationships between features (e.g., genes) and deriving low-dimensional, biologically meaningful cell embeddings^1–3^. These embeddings map phenotypically similar cells close to each other in latent space, forming the foundation for key downstream analyses such as cell annotation^4,5^, batch correction^6–8^, and multi-omics integration^2,9^.

A fundamental step in single-cell omics analysis is cell annotation, which is essential for understanding cellular heterogeneity within samples and for conducting key downstream analyses, such as trajectory inference and cell-cell communication. Modern cell annotation approaches increasingly utilize cell embeddings, followed by either clustering (marker-based) or classification (reference-based) techniques^10^. Marker-based (or cluster-centric) methods depend on well-defined clustering solutions to identify meaningful marker features. However, these clustering solutions can vary significantly depending on the user-defined clustering resolution and the chosen algorithm, potentially leading to inconsistent or inaccurate biological annotations^11^. Moreover, identifying cell types based on marker features is often a manual, labor-intensive process that requires expert knowledge^12,13^. While efforts to semi-automate this process using generative models, like GPTCelltype^13^, have shown promise, the reliance on predefined clusters and marker genes remains a limitation and a source of inconsistency.

In contrast, reference-based methods use supervised machine learning or clustering techniques to transfer cell-type labels from a reference or source dataset to a query or target dataset, resulting in automated annotation, improved accuracy, and broader applicability^7^. With the continuous expansion of single-cell sequencing datasets and the availability of annotated datasets, reference-based methods have become the preferred approach for cell-type annotation, leading to the development of diverse methods, including an increasing number of deep-learning based approaches^2,10,14^.

Despite the advantages of reference-based methods, challenges still need to be addressed. These include imbalances in cell numbers which pose challenges in identifying rare cell types, and the inability to detect novel or unknown cell types not present in the reference dataset^15^. Moreover, underlying batch effects between reference and query data are often overlooked, hindering accurate label transfer across heterogeneous datasets. Additionally, the ability to transfer labels across multi-omics modalities remains to be established, highlighting critical areas for further investigation.

To address these limitations, we propose an innovative solution: SAFAARI (<u>**S**</u>ingle-cell <u>**A**</u>nnotation and <u>**F**</u>usion with <u>**A**</u>dversarial Open-Set Domain <u>**A**</u>daptation <u>**R**</u>eliable for Data <u>**I**</u>ntegration). SAFAARI can learn domain-invariant embedding and transfer labels in the presence of batch effects, biological domain shifts, and across diverse omics modalities using an adversarial domain adaptation strategy^16^. SAFAARI can identify novel cells not present in the reference dataset through Positive-Unlabeled Learning^17^ and uses the synthetic minority oversampling technique (SMOTE)^18^ to mitigate class imbalance, enabling the annotation of rare cell types. Additionally, SAFAARI incorporates a contrastive learning strategy^19^ making it capable of integrating datasets with diverse types of batch effects or across different omics modalities. Notably, SAFFARI leverages cell type annotations through a supervised contrastive learning strategy^20^ to learn integrated joint representations across different batches or omics modalities. This approach offers a key advantage over previous methods, which are often unsupervised or tend to overlook label information, thereby improving the accuracy and consistency of cross-batch and multi-omics integration across labelled datasets.

We evaluated SAFAARI against different cell type annotation and integration methods across diverse datasets with technical batch effects and biological domain shifts, as well as simulation data and single-cell multi-omics data. Accordingly, we corroborated the following capabilities: 1) automated cell annotation via label transfer from an annotated reference or source dataset to an unannotated target dataset, 2) identification of novel cell types in the target dataset, 3) adjustment for class imbalance and detection of rare cell types via augmentation, 4) removal of batch effects, 5) integration of data sources across and within omics modalities, 6) leveraging annotations when available to enhance integration performance, and 7) scalability to large datasets.

## Results

### Overview of SAFAARI

SAFAARI is designed to perform label transfer from an annotated dataset or domain (referred to as the *source* or *reference* dataset) to an unannotated (referred to as the *query* or *target* dataset). These two domains are often heterogeneous, with different underlying distributions, commonly referred to as domain shift or batch effects. These shifts reflect unwanted technical variation (e.g., differences in protocols, experimental conditions, and sample acquisition) or biological differences (e.g., tissue types, spatial locations, species or inter-individual variability) between the source and target domains.

Illustrated in Figure 1, SAFAARI is a feedforward artificial neural network consisting of fully connected layers with nonlinear activation functions, which maps source and target cells into a shared low-dimensional latent space through representation learning^21^. This latent feature space, referred to as cell embedding, should represent similar cell types from different domains with similar embeddings.

**Figure 1.**
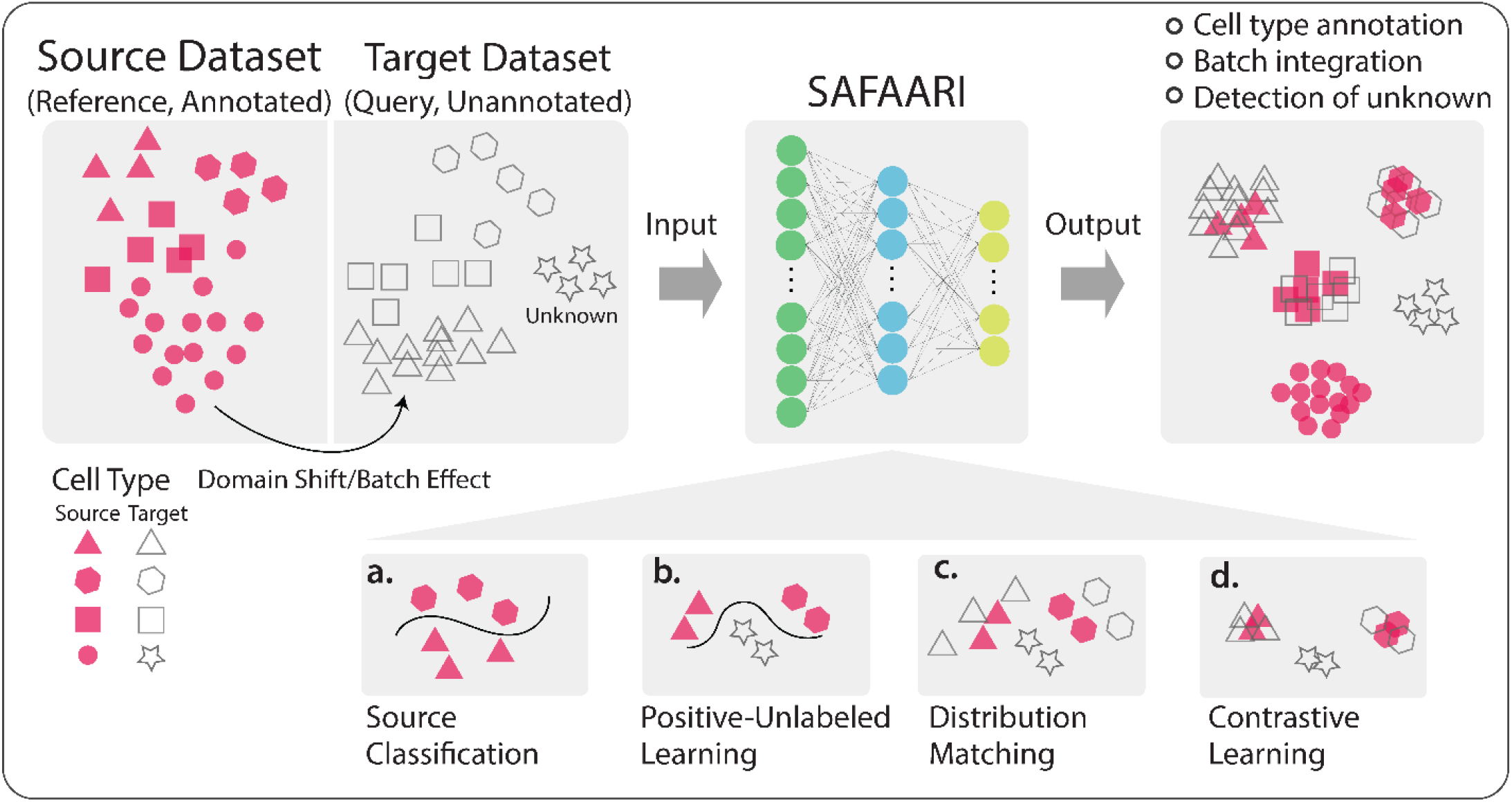
Overview of SAFAARI framework. SAFAARI is an artificial neural network that maps source and target cells into a shared low-dimensional latent space through representation learning. It optimizes a novel composite objective function that integrates four loss functions to simultaneously perform: (a) source classification, (b) positive-unlabeled learning, (c) distribution matching, and (d) contrastive learning.

To achieve this, SAFAARI optimizes a novel composite objective function (detailed in Methods), which integrates four loss functions to simultaneously conduct: **a) source classification** by training a nonlinear model on labeled source dataset to separate different cell types, **b) positive-unlabeled learning**^17^ to identify novel cell types in the target domain by assigning pseudo-labeled target cells with low-confidence predictions to an “unknown” class, **c) distribution matching** via adversarial domain adaptation^16,17^ to ensure that the cell embeddings are domain-invariant by aligning the distributions of the source and target data in the latent space, reducing the impact of domain shift, and **d) contrastive learning**^19^ to enhance the separation between different cell types and support batch integration by mixing batches within the same cell types, ultimately improving performance in batch effect correction.

### Cell type annotation in the presence of technological domain shift

Single-cell transcriptome sequencing data is generated using a variety of protocols and technological platforms, each with differences in throughput, cost, coverage, and sensitivity^22^. These methods range from microfluidic droplet-based platforms (such as 10x Genomics Chromium, Drop-seq, and inDrops) to plate-based scRNA-seq technologies like Smart-seq, Smart-seq2, and Smart-seq3, resulting in substantial heterogeneity across datasets. This variability poses challenges for label transfer in identical biological contexts, as domain shifts or batch effects are introduced by these differences in technology.

To evaluate SAFAARI’s label transfer ability in the presence of technological batch effects or domain shifts across different tissues, we used the Tabula Muris cell atlas, a compendium of single-cell transcriptome data from various tissues and organs of the model organism Mus musculus^23^. The dataset uses two distinct technical approaches for most organs: FACS-based cell captures in plates (FACS) and microfluidic droplet-based capture using 10x Genomics (10x). We focused on eight organs/tissues (bladder, kidney, heart, mammary gland, muscle, liver, bone marrow, and spleen) and preprocessed the data (see Methods) for both technologies, using either 10x or FACS as the source and target domains. Two scenarios were considered: 1) **closed-set scenario** where the same cell types were present in both the source and target domains, and 2) **open-set scenario** where a new cell type was present in the target domain but absent from the source domain (as detailed in Supplementary Table 1).

We compared SAFAARI’s average accuracy in cell type annotation with seven competing methods, including three conventional machine learning models: support vector machine (SVM)^24^ with a linear kernel, Random Forest (RF)^25^, and multilayer perceptron (MLP)^26^. Previous benchmark studies^27^ have recommended these models for cell-type identification in scRNA-seq data. Additionally, we compared SAFAARI with Seurat V4^28^, SingleR ^29^, ItClust^15^ and Concerto^2^. Seurat and SingleR were included due to their prevalence and proven performance in cell-type annotation. ItClust, to the best of our knowledge, is the only method preceding SAFAARI that claims the capability to identify clusters of unknown cells in the target dataset. However, it is constrained by a predetermined number of clusters identical in both the source and target domains. Concerto was included as the state-of-the-art benchmark method due to its recent development and the performance reported in the original study. Together, these methods provide a comprehensive benchmarking set, encompassing a range of approaches to evaluate SAFAARI’s performance thoroughly.

Our results demonstrate that SAFAARI performs as well as or better than competing methods in terms of average class accuracy in the closed-set scenario and significantly outperforms them in the open-set scenario (p-value < 4.52E-04, t-test), as shown in Figure 2A. When using 10x as the source to annotate FACS data, SAFAARI, with the exception of bone marrow, accurately identified new cell types in the target domain as “unknown.” The unknown class accuracy rates ranged from 0.31 in the heart to 0.97, 0.99, and 1.00 in the bladder, mammary, and liver, respectively, as shown in the confusion matrices in Figure 2B. Similarly, when transferring cell type annotations from FACS to 10x, SAFAARI demonstrated high accuracy across all tissues, with unknown class accuracy ranging from 0.74 in the kidney to 0.97 and 1.00 for muscle and bladder, respectively (Supplementary Figure 1).

**Figure 2.**
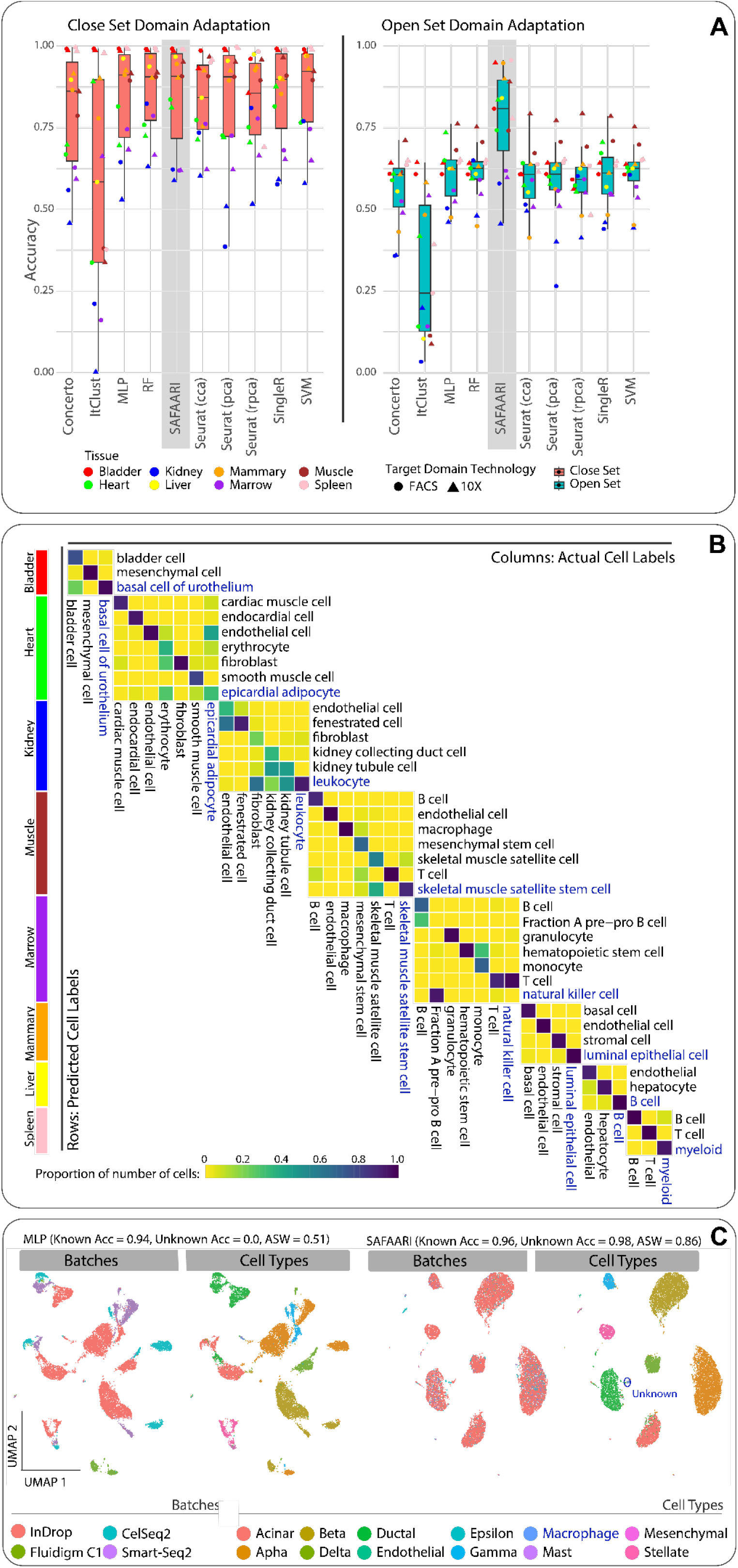
Cell type annotation in the presence of technological domain shift. **A)** Comparison of SAFAARI’s performance with the selected reference-based cell-type annotation models in both open-set and closed-set settings. The scRNA-seq data from eight different tissues in the Tabula Muris cell atlas was obtained where the gene counts were derived using two techniques: 10x Genomics and FACS-based cell capture in plates (FACS). For the performance assessment, either FACS or 10x was considered as the source dataset, and the other as the target dataset, to evaluate reference-based cell type annotation or label transfer in the presence of a technology-based domain-shift or batch effect. Two scenarios were considered: the closed-set, where only cell types common to both source and target datasets were included, and the open-set, where the target dataset contained an unknown cell type not present in the source dataset (Supplementary Table 1). **B)** Heatmap representing the confusion matrix across eight tissues (target: FACS), showing cell-type-specific annotation performance. Columns represent the actual cell labels, while rows show the predicted cell labels. The cell type coloured in navy blue represents the unknown cell type whose instances were removed from the source dataset. Colours in the viridis palette and indicate the proportion of cells relative to the sum of the column (i.e., values across columns should add up to 1.0). This represents the proportion of correct classifications (diagonal values) and misclassifications for each particular cell type represented by the column names. **C)** UMAP of open-set Label transfer result of SAFAARI on four human pancreas datasets generated with different technologies, including microfluidic (Fluidigm C), droplet-based (InDrops) and plate-based scRNA-seq (CEL-seq2, Smart-seq2) as detailed in Supplementary Table 2. It demonstrates SAFAARI’s superior batch mixing, cell separation and unknown cell type detection.

Furthermore, SAFAARI can integrate multiple source datasets to enhance the richness of the reference domain for label transfer. We tested this by collecting four human pancreas datasets generated with different technologies, including microfluidic (Fluidigm C1^30^), droplet-based (InDrops^31^), and plate-based scRNA-seq (CEL-seq2^32^, Smart-seq2^33^), as detailed in Supplementary Table 2. SAFAARI effectively annotated the cell types in the target domain (InDrops), achieving an average accuracy of 0.96 for known cell types and 0.98 for unknown cells. Furthermore, SAFAARI showed superior batch mixing and cell separation with an average silhouette width (ASW_CellType) of 0.86 (compared to 0.51 for the raw data), reflecting its ability to achieve efficient batch integration simultaneously, cell type separation and novel cell identification (Figure 1C).

### Technological batch correction and bio-conservation

Similarly, we used the Tabula Muris datasets (Supplementary Table 1) to evaluate SAFAARI’s performance in integrating technological batches. We assessed the batch integration capability of SAFAARI in comparison with six methods, including Seurat (cca and rpca)^34^, Harmony^35^, Scanorama^36^, BBKNN^37^ and Concerto^2^. These methods were selected to cover a broad range of techniques, reflecting their recent advancements (Concerto), proven performance based on prior benchmarks^38^ (Harmony), widespread use (Seurat) and intuitive approaches (BBKNN and Scanorama), providing a comprehensive evaluation framework.

The integration was evaluated using two categories of performance metrics: 1) batch effect removal (or batch mixing) and 2) conservation of biological variance from cell identity labels. For batch effect removal, we used the average silhouette width across batches (ASW_Batch), and for biological conservation, we used ASW across cell types (ASW_CellType). The ASW evaluates the cohesion within a cluster relative to its separation from the nearest neighboring cluster, with values scaled between 0 and 1. A higher ASW indicates more clearly defined clusters. When assessing batch mixing, lower ASW values are preferred, as they indicate successful integration of batches. For consistency across both metrics, we subtracted ASW_Batch from one so that higher values indicate better performance for both measures.

As shown in Figure 3A, SAFAARI consistently demonstrated the highest biological conservation, as measured by ASW_CellType, outperforming all benchmarked methods (p-value < 1.9 × 10−^3^). However, batch mixing values were generally lower across all methods. Although these methods often visually align batches (Figure 3C and Supplementary Figure 2), they do not achieve homogeneous mixing. As a result, ASW_Batch, which calculates the average relative distances between neighboring clusters, does not show significant improvement compared to the raw data, where batches remain visually distinct. This is a limitation of the metric and is consistent with previous reports^39^. Nevertheless, ASW_Batch values for all methods remained within a similar range, typically between 0.4 and 0.5, with Seurat frequently on the higher end. Despite this, qualitative evaluations based on 2D UMAP visualizations (Figure 3C and Supplementary Figure 2) indicated competitive batch alignment in SAFAARI compared to the benchmarked methods.

**Figure 3.**
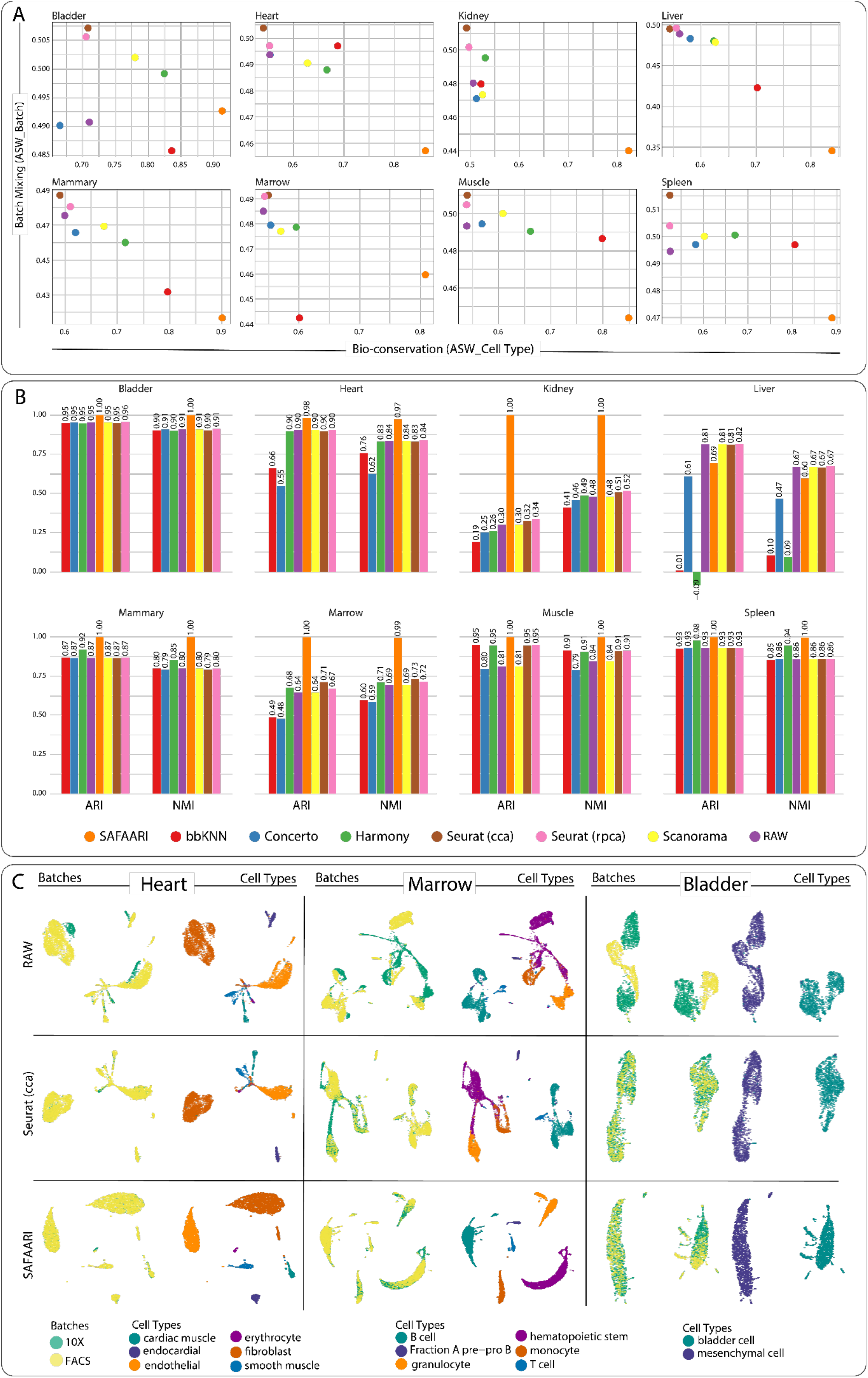
Technological batch correction and bio-conservation. **A)** Comparison of SAFAARI’s cross-technology integration performance with other integration methods. Single-cell RNA sequencing (scRNA-seq) data from eight different tissues in the Tabula Muris cell atlas were obtained, where gene counts were generated using two techniques: 10x Genomics and FACS-based cell capture in plates (FACS). For performance evaluation, we used ASW_Batch as the batch mixing metric and ASW_CellType as the biological conservation metric. **B)** Bar plots showing the comparison of SAFAARI’s clustering accuracy for different tissues with other methods, assessed by Adjusted Rand Index (ARI) and normalized mutual information (NMI). **C)** UMAP visualizations of SAFAARI’s integration for Heart, Marrow, and Bladder. UMAPs of other tissues are presented in Supplementary Figure 2.

We also evaluated the clustering accuracy of cell types in the integrated datasets using clustering, assessed by Adjusted Rand Index (ARI) and normalized mutual information (NMI) for global cluster matching (Methods). SAFAARI consistently achieved the highest ARI and NMI scores across all tissues, except for the liver, which still returned a competitive performance (Figure 3B).

These experiments highlight the utility of supervised integration when labels are available. Nonetheless, SAFAARI can also effectively integrate an unannotated query dataset into a reference dataset using pseudo-labels identified through label transfer (Supplementary Figure 3).

### Multi-batch integration in simulation datasets

To evaluate integration methods in a controlled setting where the nature of the batch effects and ground truth are known, we used two simulated datasets (Supplementary Table 3) with varying complexities, including different numbers of batches, cell types within batches, cells, and scaling factors affecting batch variation^40^.

Figure 4 presents UMAP plots showing batches and cell types in both the unintegrated data and after integration using SAFAARI and three other competing methods, selected based on their performance in our previous evaluations. Simulation task 1 posed a higher level of complexity, and therefore, integration techniques often struggle to effectively mix batches while maintaining separation between cell types, as highlighted in prior benchmarking studies covering a broad range of methods^38^. SAFAARI demonstrated an apparent ability to separate cell types while successfully integrating batches in both simulation tasks. Notably, in the more complex Simulation Task 1 (Figure 4A), where other methods struggled, SAFAARI excelled at integrating the data while maintaining cell type separation.

**Figure 4.**
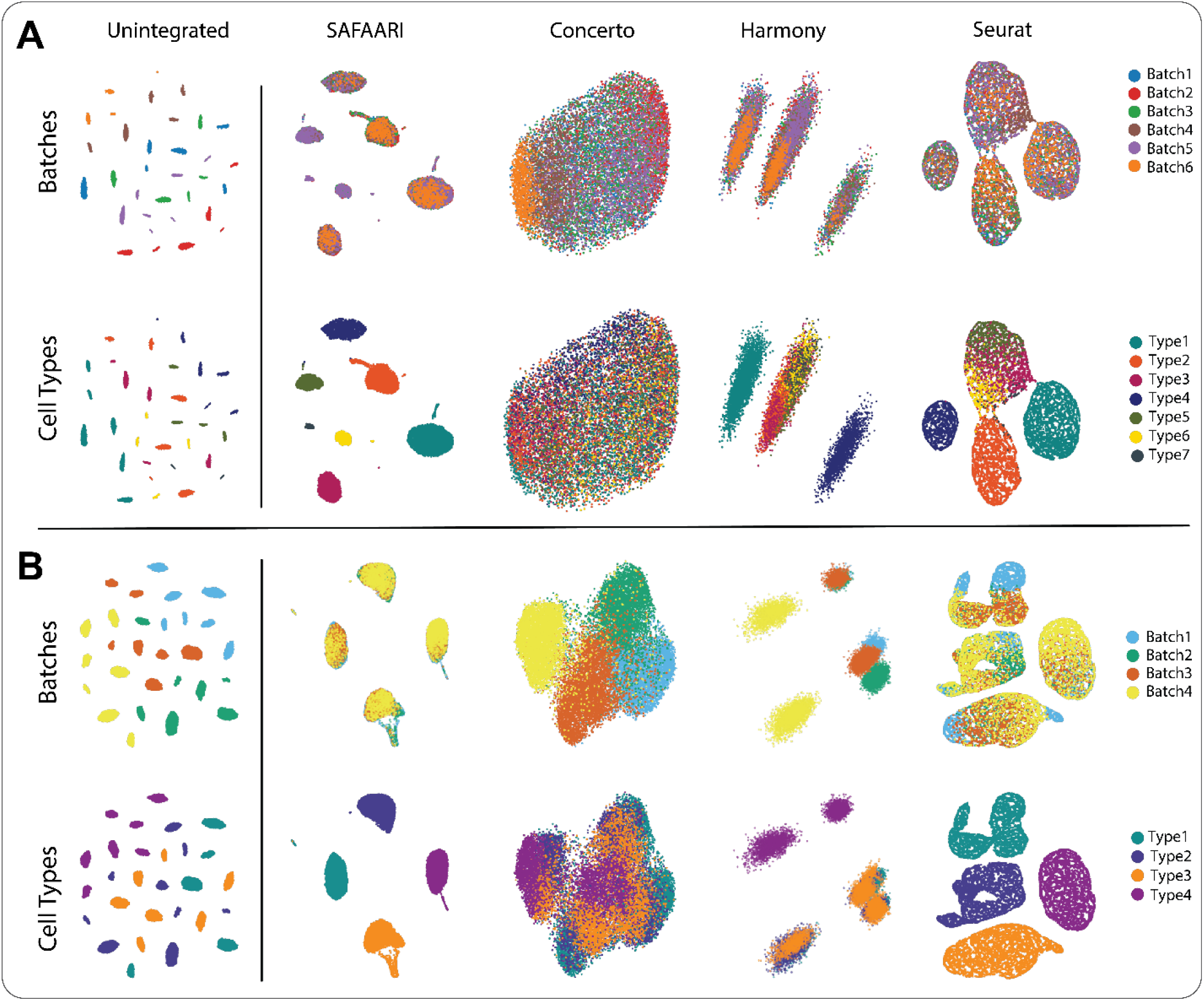
Multi-batch integration in simulation datasets. **A)** UMAP plots illustrating batches and cell types in both unintegrated data and after integration using SAFAARI, along with two competing methods for the complex simulation dataset 1 (Supplementary Table 3). **B)** UMAP plots showing batches and cell types in both unintegrated data and after integration using SAFAARI and two competing methods for simulation dataset 2 (Supplementary Table 3).

### Cell type annotation in the presence of biological domain shift

We retrieved single-cell expression data from the Human Lung Atlas^38^, which includes samples from three different laboratories, generated using Drop-seq and 10x Chromium platforms, and comprising both transplant and biopsy data. The lung biopsy samples were obtained from a distinct spatial location (airways) compared to the transplant samples (parenchyma). We used the transplant dataset, consisting of 22,417 cells across 14 cell types, as the reference to annotate the biopsy dataset, which contained 4,998 cells across eight cell types, including Ionocytes, a rare cell type with 46 cells not present in the transplant (reference) dataset (Supplementary Table 4). The labels for the target biopsy dataset were blinded and input into SAFAARI for annotation. This task presented a significant challenge due to multiple layers of domain shift, including differences in technologies, laboratory origins, individuals, and spatial sampling locations. Moreover, the cellular composition varied significantly between the datasets: in the biopsy data, ciliated and secretory cells accounted for over 86% of the total, while in the transplant dataset, these cells comprised less than 6%, with immune cells being predominant (Supplementary Table 4).

Despite these complexities, SAFAARI achieved a mean accuracy of 0.85 for shared cell types and 0.70 for the novel cell type, correctly identifying 32 of the 46 Ionocyte cells as “unknown” (Figure 5A). Additionally, immune cells, although underrepresented in the biopsy data compared to the transplant reference, were generally predicted correctly or misclassified as another immune cell type, which is phenotypically more similar than ciliated, secretory, or endothelial cells, providing a biologically plausible explanation for these misclassifications.

**Figure 5.**
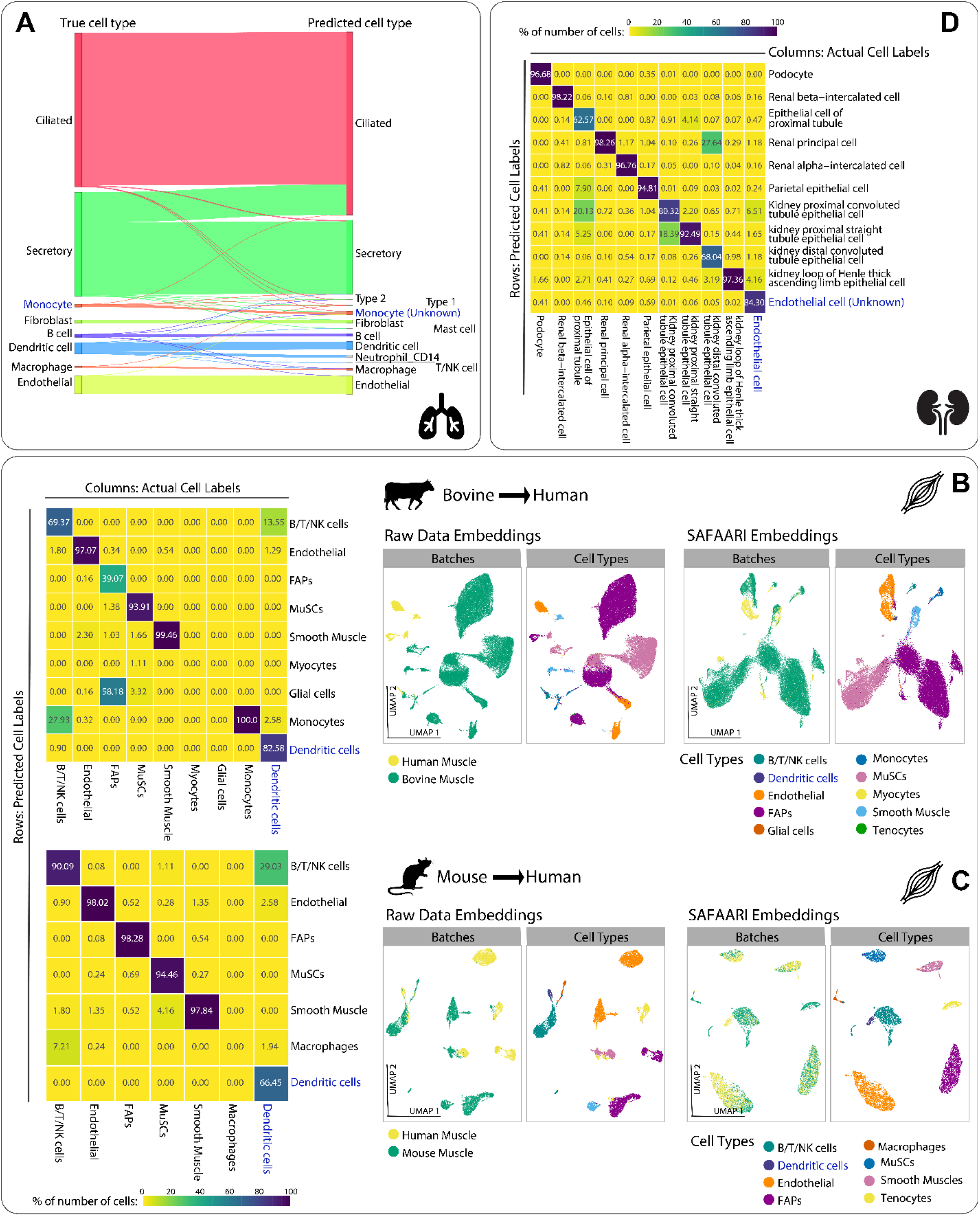
Cell type annotation in the presence of biological domain shift. **A)** Sankey plot showing SAFAARI’s open-set domain adaptation on Human Lung Atlas samples from three laboratories, using Drop-seq and 10x Chromium platforms, covering both transplant and biopsy data (Supplementary Table 4). The transplant dataset served as the reference to annotate the biopsy dataset, including rare Ionocytes absent in the reference. Labels for the biopsy dataset were blinded and annotated using SAFAARI. **B)** UMAP visualization and heatmap representing the confusion matrix for cross-species open-set domain adaptation results on scRNA-seq muscle data, used to annotate cell types in the human dataset with bovine as the source domain (Supplementary Table 5). Columns represent actual cell labels, while rows show predicted cell labels. The cell type colored in navy blue represents the unknown cell type, whose instances were removed from the source dataset. Colors follow the viridis palette and indicate the proportion of cells relative to the sum of the column (i.e., values across columns sum to 1.0). This illustrates the proportion of correct classifications (diagonal values) and misclassifications for each cell type represented by the column names. **C)** UMAP visualization and heatmap representing the confusion matrix for cross-species open-set domain adaptation results on scRNA-seq muscle data, used to annotate cell types in humans with mouse as the source domain (Supplementary Table 5). **D)** Heatmap representing the confusion matrix for open-set domain adaptation results on single-nucleus ATAC-seq data from two distinct studies of adult human kidney cortex samples (Supplementary Table 6).

We also explored cross-species domain adaptation, one of the most challenging tasks due to the substantial biological differences between species. However, such comparative analysis holds great potential for uncovering fundamental evolutionary processes, such as identifying conserved cell types across species and the gene programs driving their similarities and differences^8^. To evaluate SAFAARI’s ability to adapt across species, we extracted scRNA-seq muscle data from three species: mouse (4,414 cells, 7 types)^41^, bovine (36,129 cells, 11 types)^42^, and human (2,876 cells, 7 types)^43^ (Supplementary Table 5). We performed two separate analyses to transfer labels from bovine and mouse (reference) to human (target).

Similar to previous experiments, the cell composition varied significantly between the reference and target datasets. In humans, endothelial cells comprised 44% of the population, while in bovine and mouse, FAPs (fibro/adipogenic progenitor cells) represented the majority (59% and 28%, respectively). For both crossspecies transfers, dendritic cells, which were present in the human dataset (n=155) but absent in the mouse dataset, were treated as an “unknown” cell type (synthetically removed from the bovine dataset as well).

After blinding the human cell labels, SAFAARI was applied to annotate human cells using the bovine reference, achieving an average accuracy of 0.83 across shared cell types and 0.82 for the unknown cell type (Figure 5B). Similarly, when annotating human cells using the mouse reference, SAFAARI reached

0.91 average accuracy on shared cell types and 0.66 on the unknown cell type (Figure 5C). Moreover, monocytes, representing a rare population in the human dataset (n=37), were synthetically removed from the reference datasets, and the experiments were repeated. However, SAFAARI was unable to predict monocytes as an unknown cell type (Supplementary Figure 4), instead mapping them to macrophages (mouse as reference) or dendritic cells (bovine as reference).

Beyond scRNA-seq data, we evaluated SAFAARI’s open-set domain adaptation capability to analyze cellular heterogeneity based on cell-type-specific chromatin accessibility profiles. Specifically, we utilized single-nucleus ATAC-seq data from two distinct studies of adult human kidney cortex samples: 1) Muto et al.^44^, which provided normal kidney tissue (non-tumor samples from patients undergoing nephrectomy), and 2) Wilson et al.^45^, where normal samples were obtained from patients undergoing nephrectomy and deceased organ donors.

To assess SAFAARI’s performance, we transferred labels from Muto et al.^44^ to Wilson et al.^45^, designating endothelial cells (n = 1,274, representing 3.5% of the target domain) as an “unknown” cell type due to their relatively distinct phenotype compared to epithelial and β-/α-intercalated cells, which dominate the shared cell types (Supplementary Table 6). SAFAARI achieved an accuracy of 0.89 for shared classes and 0.84 for the unknown class (Figure 5C). The lowest per-class accuracy was observed for ‘epithelial cell of the proximal tubule’ (accuracy = 0.63), where 20% of misclassifications were assigned to ‘kidney proximal convoluted tubule epithelial cells’, both represent epithelial cells of the proximal tubule, different based on their location in the cortex and the course of the tubule (convoluted vs. straight). In a subsequent experiment, we considered T cells—a rare cell type (n = 82, representing 0.2% of the target domain)—as the unknown class and repeated the analysis. SAFAARI produced comparable results, with an average accuracy of 0.88 for shared cell types and 0.76 for identifying T cells as unknown (Supplementary Figure 5).

### Multi-omics integration

Advancements in single-cell sequencing technologies have produced extensive multi-modal omics data, particularly in single-cell RNA and ATAC sequencing. Joint analysis of these datasets has greatly improved our understanding of cellular heterogeneity and regulatory networks, establishing multi-omics integration as a critical step in single-cell research^46^. This integration can be categorized into matched or unmatched pipelines: matched pipelines measure multiple omics layers from the same cell, while unmatched pipelines involve unaligned layers generated from separate experiments^47^.

Accordingly, we evaluated SAFAARI’s ability to integrate transcriptome and chromatin accessibility data using two datasets: 1) a molecular atlas of the normal postmenopausal ovary (unmatched), comprising 44,402 cells across both modalities and five cell types^48^ (Supplementary Table 7) and 2) a 10x multiome dataset of peripheral blood mononuclear cells (PBMCs) (matched), containing 10,412 cells and 19 distinct immune cell types^49^ (Supplementary Table 8). In both datasets, SAFAARI effectively integrated scRNA-seq and scATAC-seq batches, as measured by ASW_Batch, while preserving biological distinctions among cell types, evaluated by ASW_CellType and clustering accuracy (NMI and ARI), as shown in Figure 6A-B. SAFAARI outperformed Harmony and Seurat (Supplementary Figure 6), as evidenced by improved quantitative integration metrics and enhanced visual inspection, demonstrating the advantages of supervised batch correction leveraging labels when available, compared to previous unsupervised methods.

**Figure 6.**
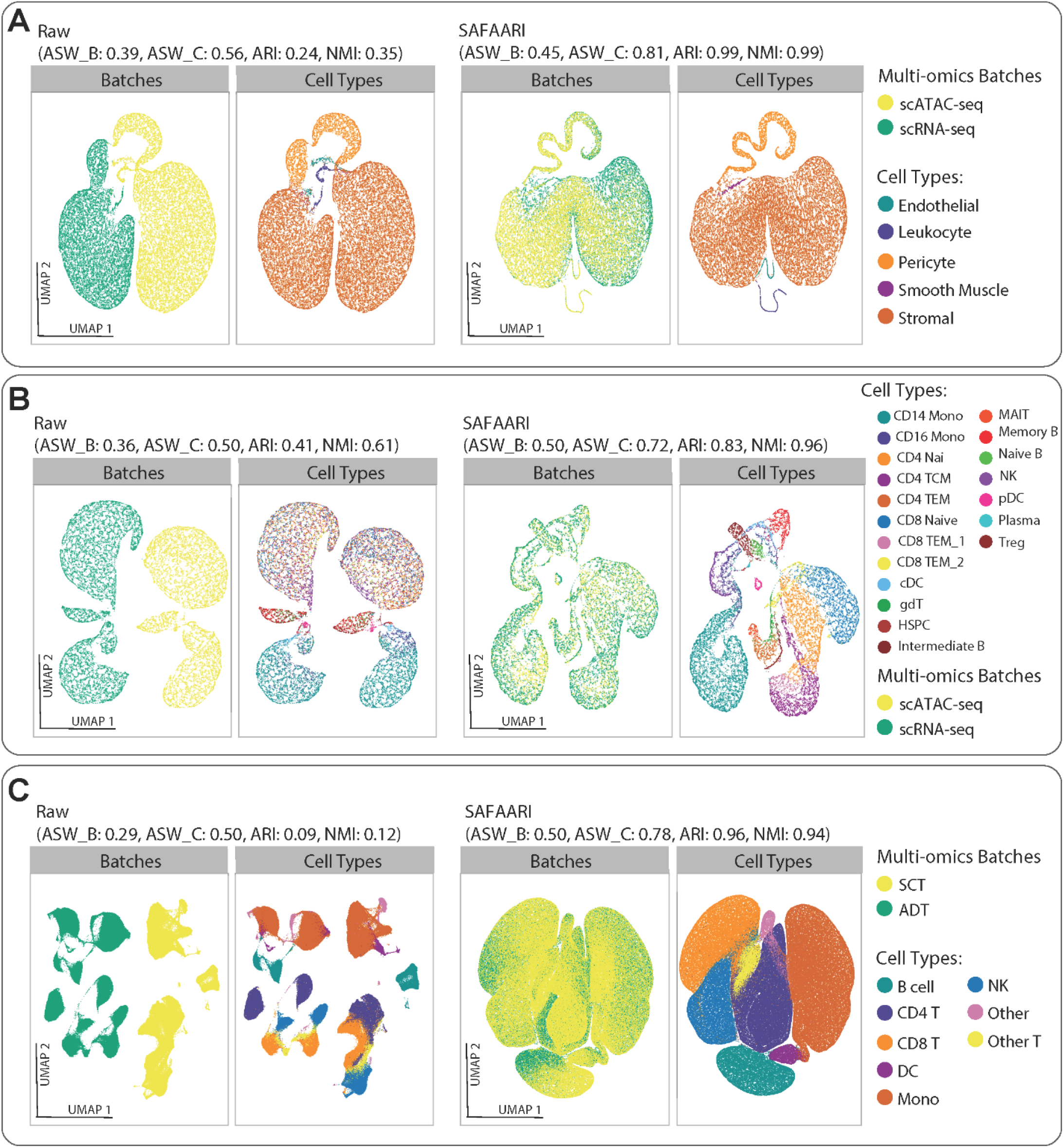
Multi-omics integration. UMAP visualization and integration metrics, including ASW_Batch, ASW_CellType, NMI, and ARI, for: **(A)** SAFAARI’s integration of scRNA-seq and scATAC-seq modality batches from a molecular atlas of the normal postmenopausal ovary (Supplementary Table 7), **(B)** SAFAARI’s integration of scRNA-seq and scATAC-seq modality batches from a 10x multiome dataset of peripheral blood mononuclear cells (PBMCs) (Supplementary Table 8), and **(C)** SAFAARI’s integration of scRNA-seq and scATAC-seq modality batches from a large-scale CITE-seq dataset of human PBMCs (Supplementary Table 9).

To further showcase multi-omics integration across different modalities, we analyzed a large-scale CITE-seq dataset^34^ of 162,000 human PBMCs, which measures both RNA and protein expression simultaneously using RNA sequencing combined with antibody-based protein labeling(Supplementary Table 9). SAFAARI effectively corrected the multi-modal batches while maintaining a clear separation of cell types, as illustrated in Figure 6B.

## Discussion

SAFAARI presents a comprehensive platform for single-cell annotation, multi-omics integration, batch correction, and visualization in single-cell sequencing data. Its innovative and flexible framework optimizes multiple loss functions simultaneously, enabling cell type annotation (classification loss), batch correction (adversarial loss), detection of unknown cell types (positive-unlabeled loss), and improved cell alignment and biological conservation (contrastive loss) across batches and omics modalities. SAFAARI’s approach to multi-omics fusion is adaptable to both matched and unmatched multi-omics datasets by employing nonlinear joint manifold learning to align different omics layers, with the potential of reducing inter-modality information loss^50^.

We demonstrated SAFAARI’s performance in label transfer across technological and biological domain shifts, integration of multi-batch real and simulated data, and across various omics modalities, comparing its efficacy to competing methods. SAFAARI pioneers a methodology centered on optimizing an objective function, offering flexibility in model architecture. Unlike previous approaches emphasizing specific model designs, SAFAARI can employ various architectures—ranging from multi-layer perceptron to transformers—to optimize its combined objective function for learning low-dimensional cell embeddings through joint representation learning. This framework allows for ongoing innovation and improvements to SAFAARI with diverse architectural advancements.

Implemented on GPU, SAFAARI is highly scalable and capable of handling large single-cell datasets efficiently. Notably, it is the first tool capable of detecting unknown cell types within a query dataset that may have an arbitrary number of cell types in either source/reference or target datasets. However, while effective, SAFAARI currently classifies all novel cells as a single “unknown” category, which limits its capacity to distinguish between diverse new cell types and reduces its predictive power. As a future enhancement, we aim to integrate a Self-supervised Class-Discovering Adapter^51^ to automatically estimate and categorize unknown cells into distinct classes, thereby improving prediction accuracy. Additionally, SAFAARI can be extended to multi-adversarial domain adaptation^52^ enabling simultaneous integration of multiple datasets or omics modalities even in the presence of class imbalance—overcoming a key limitation of existing methods that are typically restricted to label transfer between two datasets or sequential dataset integration.

Overall, SAFAARI offers an innovative, robust, accurate, scalable, and flexible platform for single-cell data analysis, suitable for a wide range of applications and biological or clinical research questions.

## Methods

### The SAFAARI model

SAFAARI is designed to identify cell types in a query or target domain by leveraging latent knowledge from a related source or reference domain. Unlike traditional label transfer methods, SAFAARI can detect novel cell types in the target domain that do not appear in the source domain. The model provides accurate cell type annotations for known classes or cell types (those present in the source domain) and categorizes unrecognized cells as “unknown” when they do not correspond to any class in the source domain.

The model employs the synthetic minority oversampling technique (SMOTE)^18^ on all labeled source samples to generate a balanced source dataset for downstream analysis. This approach ensures that the number of samples in each class matches the count of the largest class in the original dataset, thereby facilitating the annotation of rare cell types.

SAFAARI’s loss functions integrate four objective functions to simultaneously optimize for domain adaptation, cell type annotation, novel cell type identification, and the separation of cell types while integrating batches across experiments. The formulation of the objective functions for SAFAARI is detailed below.

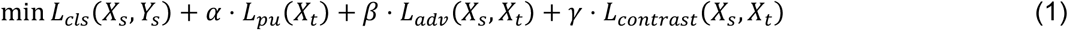

X_*s*_ rerefers to the input data representing the omics profiles of cells in the source domain, while *Y*_*s*_ denotes the corresponding ground-truth labels for X_*s*_.Similarly, X_*t*_ represents the input data from the target domain.

First, a classifier is trained on the cell types from the source domain, where *L*_*cls*_ (X_*s*_, *Y*_*s*_) represents the classification loss over this reference domain. However, since the target domain may contain unknown cell types, a classifier trained solely on *L*_*cls*_(X_*s*_, *Y*_*s*_) cannot distinguish between known and unknown cell types in the target domain. To overcome this limitation, we introduce a positive-unlabeled learning^53^ loss *L*_*pu*_(X_*t*_), following authors’ previous work^17^, to identify unknown samples in the target domain.

Furthermore, to minimize distributional shifts between the source and target domains, we apply an adversarial loss *L*_*adv*_ (X_*s*_, X_*t*_), which aligns their distributions within a shared latent embedding space. Additionally, a supervised contrastive learning loss, *L*_*contrast*_ (X_*s*_, X_*t*_), is used to enhance separation between samples from different cell types while mixing the batches of the same cell types.

The objective function of SAFAARI is motivated by the theoretical analysis of open-set domain adaptation presented in our previous work^17^, which demonstrates that the prediction error on the target domain is bounded by the prediction error on the source domain, the positive-unlabeled learning risk on the unlabeled target domain, and the distributional discrepancy between the source and target domains. Thus, SAFAARI aims to minimize this upper bound on the prediction error in the target domain. The following sections describe SAFAARI’s architecture and detail each component of its objective function.

### Feature Embedding

The first step in the SAFAARI framework is to map both reference and target samples into a shared embedding space using a neural network. Specifically, we employ a two-layer multi-layer perceptron (MLP) with 168 and 32 hidden neurons in the respective layers to learn the features of both source and target samples. For each source cell 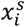 and target cell 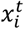, the MLP is applied to extract the feature representations, as follows:

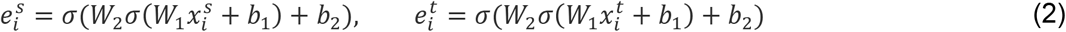

where, *W*_1_ and *W*_2_ represent the trainable weight parameters, *b*_1_ and *b*_2_ denote the respective trainable bias parameters and *σ*(·), denotes a non-linear activation function, i.e., ReLU.

### Classification loss

The first term, *L*_*cls*_ (X_*s*_, *Y*_*s*_), in the SAFAARI objective function, is designed to learn the class memberships from the source training samples. It is defined as follows:

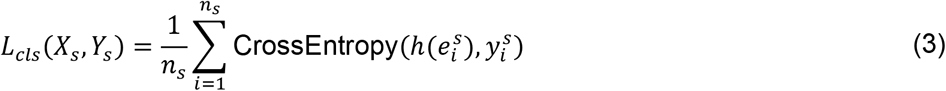

Here, CrossEntropy(.) denotes the cross-entropy loss, and *h*(.) is a linear classifier:

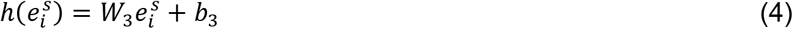

### Positive-unlabeled loss

The second term, *L*_*pu*_(X_*t*_) of SAFAARI loss function is inspired by positive-unlabeled learning^17,53,54^. Originally, positive-unlabeled learning^53^ addressed a binary classification problem with only positive and unlabeled samples (which could be either positive or negative). The goal is to train a classifier to distinguish between positive and negative samples. More recently, this approach has been extended to open-set domain adaptation scenarios^17^, where the reference data from the known cell types are treated as positive samples, and the target data are considered unlabeled samples. In this context, the objective of positive-unlabeled learning can be expressed as follows:

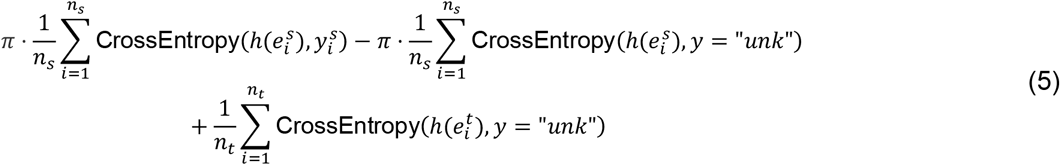

Here, *π* = 1 − *p*(*y* = “*unk*”), where *p*(*y* = “*unk*”) denotes the class-prior probability of unknown samples in the target domain. CrossEntropy 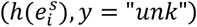 represents the cross-entropy loss when predicting 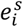 as belonging to the unknown cell types.

Under mild conditions, as demonstrated in the literature^53^, this objective can be simplified to:

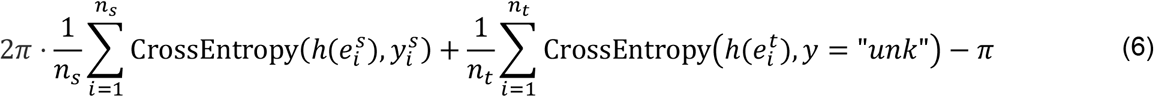

The mild conditions refer to the symmetry of the loss function, where for a given function *f*(*x*), the loss function *L* satisfies a symmetric property:

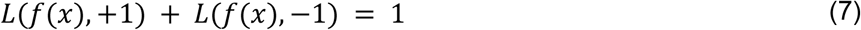

This symmetry implies that the total loss for positive and negative samples is balanced, ensuring consistent behavior for both types of samples.

However, in practice, accurately estimating the class-prior probability *π* is challenging. As a result, the second term, 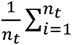 CrossEntropy 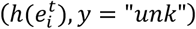, may lead to a suboptimal solution if all target samples are treated as belonging to the unknown cell type. To address this issue, we employ pseudo-labeling techniques in this paper. Specifically, the classifier *h*(·) is used to generate pseudo-labels for the target samples. We then select unknown candidate samples based on the maximum prediction probability produced by *h*(·). The underlying intuition is that if a target sample belongs to one of the known cell types, *h*(·) will assign it to the correct cell type with high confidence. Conversely, if the sample is unknown, the maximum prediction probability will be lower. Therefore, we simply select *k* target samples with the lowest maximum prediction probabilities. The first term of positive-unlabeled learning corresponds to the classifier learning, *L*_*cls*_ (X_*s*_, *Y*_*s*_). The second term contributes to the following loss term:

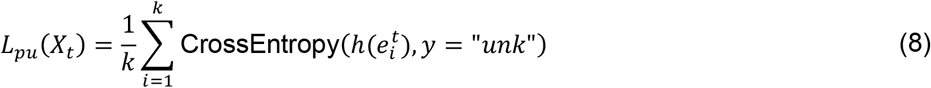

which is defined based on the selected pseudo-labeled target samples.

### Adversarial loss

The third term, *L*_*adv*_ (X_*s*_, X_*t*_), of the SAFAARI’s objective function is designed to minimize the distributional discrepancy between the reference and target domains in the new feature space. The objective of *L*_*adv*_ (X_*s*_, X_*t*_) is to learn domain-invariant representations for both reference and target samples. In this paper, we employ an adversarial learning approach^55^ to quantify the distributional discrepancy between the two domains. This approach begins by assigning domain labels to all training samples: source samples are labeled as *d* = 0, and target samples as *d* = 1. A domain discriminator:

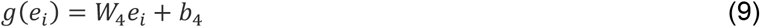

where W_4_ and, *b*_4_ are trainable parameters, is then used to differentiate between the reference and target samples.

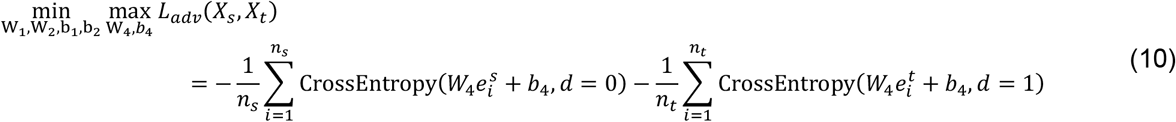

The minimax optimization of *L*_*adv*_ (X_*s*_, X_*t*_) indicates that the feature embedding function aims to learn the domain-invariant representations so that the domain discriminator *g*(·) gets confused about the domain label prediction. In contrast, the domain discriminator *g*(·) aims to predict the domain label correctly. After optimization, source and target domains will have the matched distributions in the embedding space.

### Contrastive loss

The fourth term, *L*_*contrast*_ (X_*s*_, X_*t*_) of SAFAARI loss aims to enhance the class separability of both source and target samples. This term further improves the distinction between known and unknown samples in the target domain. In this study, we employ supervised contrastive learning^20^, which constructs positive sample pairs based on cell types. For target samples, pseudo-labels generated by the classifier *h*(·) are used. Positive pairs are formed when two samples belong to the same cell type. Following the recommendation^20^, the feature embeddings are first normalized, and then the contrastive loss is computed as follows:

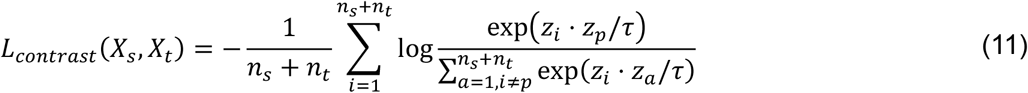

where (*z*_*i*_, *z*_*p*_) represents the positive pair and *τ* is the adjustable temperature coefficient (*τ* = 0.07 in this study).

### Data preprocessing

All data follow the same preprocessing procedure. First, for both the source and target data, a cell is filtered out if the number of genes with non-zero expression is less than 100. We further remove MT (Mitochondrial) and ERCC (External RNA Control Consortium spike-ins) genes and genes that are expressed in less than ten cells; the gene expression values are then normalized.

To ensure that gene expression levels are comparable across different cells, we employed a standard preprocessing pipeline that involves normalization, scaling, and log transformation of the data. Initially, each gene’s expression level in a cell is normalized by dividing it by the total gene expression of that cell. This step corrects for variations in sequencing depth, making the expression levels more comparable across cells. After normalization, the expression values are scaled by multiplying them by 1000,000. This scaling step, which is the Counts Per Million” (CPM) method^56^, helps to bring the data into a more interpretable range without distorting the relative expression levels.

Finally, we apply a natural logarithm transformation to the scaled values. This transformation is crucial for stabilizing variance across the data and mitigating the influence of extreme outliers, thereby making the data distribution more normal. The log transformation effectively compresses the expression level range, facilitating downstream analyses.

Highly variable genes (HVGs) are first selected from the target dataset, while a similar selection is made from the source domain. The next step involves finding the intersection of these two gene sets, and the overlapping genes are then chosen for further use in the model. We infer that this approach enhances the relevance and robustness of the analysis. Highly variable genes (HVGs) exhibit significant variability across cells, often highlighting critical biological differences, such as cell type-specific expression patterns. By intersecting HVGs from both the target and reference datasets, this method effectively filters out dataset-specific noise, ensuring that only the most informative and consistently relevant genes are utilized. This strategy improves the model’s generalizability across different datasets by capturing the unique characteristics of the target dataset while still maintaining broader applicability. Consequently, the analysis becomes more robust, reducing the risk of overfitting and enhancing the model’s performance on new data, which is particularly crucial in transfer learning scenarios.

For open-set annotation scenarios, in all datasets, any cell type present in the target dataset but absent in the reference dataset was designated as the ‘unknown’ class. For target datasets that did not contain such an unknown cell type, we synthetically removed a cell type, preferably the one with the fewest cells, to simulate the concept of rare, novel, or unknown cell types.

### Compared method for cell annotation

We evaluated SAFAARI’s annotation capability by comparing its performance against seven competing methods selected for either their widespread usage or recent standing as state-of-the-art approaches.

#### Seurat V4

Seurat is a widely used tool for single-cell RNA sequencing (scRNA-seq) data analysis among biologists and clinical experts. As part of its annotation capabilities, Seurat V4^34^ implements a reference-based annotation approach, mapping query samples to a well-annotated reference dataset. In this study, we utilized Seurat V4’s label transfer feature, which relies on the ‘*FindTransferAnchors’* function to identify anchors—pairs of cells from the reference and query datasets that share similar biological characteristics. These anchors enable the transfer of cell type labels from the reference to the query dataset through the ‘*TransferData*’ function.

While Seurat 4.0 leverages information from both the source and target datasets by identifying anchor pairs, it does not explicitly use cell type labels from the source data. An optimal method for cell type identification should incorporate cell-type-specific gene expression from both well-annotated source data and unlabeled target data. Since the source and target datasets contain varying degrees of cell-type-specific information, a data-driven approach that adjusts the contribution of each data type during analysis would be preferable.

To ensure comprehensive analysis, this process was repeated using various dimensionality reduction methods, including PCA, reciprocal PCA (RPCA), and canonical correlation analysis (CCA), allowing us to compare the effectiveness of each method in accurately annotating cell types.

#### SingleR

SingleR is a reference-based analysis method designed for cell type annotation in scRNA-seq data. It works by calculating the Spearman correlation coefficient for variable genes, which is then aggregated to score each cell against different cell types. The method iteratively refines this process by subsampling the top-scoring genes, allowing it to accurately distinguish between closely related cell types. Although SingleR is a powerful tool for initial cell type annotation, particularly when using high-quality reference datasets, its performance is highly dependent on the quality, comprehensiveness, and resolution of the reference data. If the reference dataset lacks certain cell types present in the query dataset, SingleR may fail to accurately annotate novel or rare cell types. While efficient for small to moderate datasets, SingleR may not scale as effectively for larger scRNA-seq datasets, where computational performance becomes a concern. Additionally, SingleR does not inherently correct for batch effects, unlike methods such as Seurat or Harmony, which explicitly address this issue. Consequently, when the query dataset includes cells from multiple conditions or batches, annotation accuracy may be affected unless batch effects are corrected prior to analysis.

#### ItClust

ItClust is a supervised clustering algorithm for single-cell RNA sequencing (scRNA-seq) data. It leverages transfer learning to integrate and utilize well-annotated reference datasets for accurate clustering and cell type classification in target datasets. It employs a stacked autoencoder for dimensionality reduction, initializing cluster centroids based on the reference data’s annotated cell types. ItClust iteratively updates both the neural network parameters and cluster centroids to adapt to the unique characteristics of the target data. ItClust is limited to a specific number of clusters predetermined in source domain and this could lead to misclassifications or a failure to recognize novel cell types in the target domain.

#### Concerto

Concerto predicts cell types by generating informative cell embeddings and identifying key gene features. The model clusters cells with similar gene expression profiles close together in the embedding space, often aligning these clusters with distinct cell types. Additionally, the model’s attention mechanism highlights the genes most important for distinguishing between these types. By comparing the embeddings of cells with well-annotated reference cells and considering the significant gene features identified by the model, Concerto effectively predicts the cell types of new samples.

### Compared method for batch integration

#### Seurat (V4)

Seurat integrates single-cell RNA sequencing (scRNA-seq) data from multiple sources. We utilized Seurat’s ‘*FindIntegrationAnchors’* function to identify anchor points between the datasets. These anchors represent biologically similar cells across different datasets, forming the foundation for the integration process. The identification of these anchors is guided by the specified reduction methods, such as reciprocal PCA (RPCA) and canonical correlation analysis (CCA), which facilitate deeper alignment of the datasets. Once the anchors are identified, the ‘*IntegrateData’* function is used to merge the datasets into a single, integrated Seurat object. Seurat v4 tend to favor the removal of batch effects over conservation of biological variation. In our analysis, we ran Seurat V4 using CCA and RPCA alignments (version 4.3.0.1).

#### Harmony

The Harmony algorithm starts by positioning all datasets in PCA space, incorporating their respective batch variables. It then alternates between two primary steps until convergence. First, it employs a maximum diversity clustering technique that prevents overcorrection by pushing clusters of similar cells apart. Subsequently, it corrects batch effects using a linear mixture model. This iterative process produces a corrected embedding that effectively mitigates batch effects while preserving the underlying biological variation. While Harmony is a powerful and flexible tool for batch correction, its primary focus is on removing batch effects, sometimes at the expense of preserving biological variation. As a result, it performs well in simulations and real datasets with less complex biological structures. However, in cases where biological compositions differ significantly between batches (e.g., the presence of novel cell types in one batch but not others), Harmony may struggle to accurately align cell types, potentially leading to misalignment^38^. For optimal results, careful parameter tuning is crucial, though in this study, we used the default settings.

#### Scanorama

The Scanorama algorithm is an advanced extension of the Mutual Nearest Neighbors (MNN) approach, inspired by the concept of panoramic stitching. It identifies similar cells across datasets through a k-nearest neighbor search and then refines these connections to establish mutual nearest neighbors. After this, all data points are embedded into a unified hyperplane. We executed Scanorama using the ‘*correct_scanpy’* function.

#### BBKNN

BBKNN starts by constructing a k-nearest neighbor graph within each individual batch. It then expands this by identifying the k-nearest neighbors among all cells across different batches. Since this process can introduce irrelevant connections between distinct cell types, BBKNN computes a connectivity score for each cell pair, like the approach used in the UMAP algorithm. This score is symmetrized to represent the strength of the connection between each pair of cells. Ultimately, BBKNN generates a weighted neighborhood graph that effectively captures the relationships between cells across batches. Although BBKNN is highly time- and memory-efficient, it tends to prioritize removing batch effects over preserving biological variation.

#### Concerto

Concerto is a self-distillation contrastive learning framework specifically tailored for the integration and analysis of multimodal single-cell data. It can also be used for cell type annotation. It features an asymmetric teacher-student network structure, where the teacher network utilizes an attention mechanism to aggregate gene embeddings into cell embeddings, and the student network converts input data into cell embeddings through a dense layer. The training objective of Concerto is to maximize the agreement between the teacher and student embeddings for the same cell, treating each cell as a unique instance while minimizing the similarity between different cells within the batch. This contrastive learning strategy produces robust and biologically meaningful cell embeddings. For data integration, Concerto incorporates a domain-specific batch normalization layer to align cell distributions across batches, effectively correcting batch effects while preserving biological signals. In the context of multi-omics datasets, Concerto integrates different modalities by summing their outputs to create unified cell representations.

### Performance evaluation metrics for annotations

To evaluate the performance of annotations, we calculated the mean class accuracy, given the multi-class nature of the dataset. Mean class accuracy is defined as the average ratio of correctly assigned cells to the total number of cells within each class, providing an overall measure of the model’s performance across all classes by accounting for its accuracy within each individual class. Additionally, we report the unknown class accuracy, which reflects SAFAARI’s ability to detect novel cell types in the target domain.

### Performance evaluation metric for integration

Batch integration evaluation is performed qualitatively through visual inspection using t-distributed Stochastic Neighbor Embedding (t-SNE)^57^ or Uniform Manifold Approximation and Projection (UMAP)^58^ 2D plots. While assessing integration quality in smaller datasets with few cell types is relatively straightforward, the evaluation becomes more subjective and challenging when comparing different algorithms on large, complex datasets. This difficulty is further compounded when batches are not clearly distinct or when similar cell types are intermixed. To address this, in our study, we utilized UMAP visualization alongside quantitative metrics such as average silhouette width (ASW)^59^, adjusted Rand index (ARI)^60^, and normalized mutual information (NMI)^61^ to evaluate batch correction methods, focusing on assessing batch mixing and cell type purity.

### Adjusted rand index (ARI)

ARI is used to quantify clustering accuracy and measures the similarity between two clustering results, defined as:

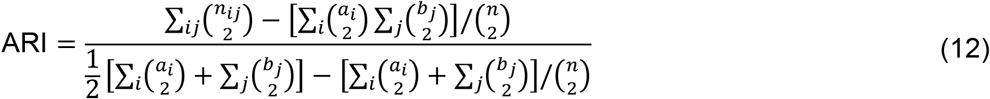

where *n*_ij_ is the number of cells in both cluster *i* of the clustering result and cell type *j* of the true cell type labels, *a*_*i*_ is the number of cells from cluster *i, b*_j_is the number of cells from cell type *j*, and *n* is the total number of cells. We calculate ARI to compare the clustering result of integrated data with the predefined cell types. ARI ranges in [0, 1], and higher values indicate higher similarities. We used the function ‘*adjusted_rand_score’* in the Python module *sklearn*.*metrics*.*cluster* implementation of the ARI. To calculate ARI, we utilized the Louvain^62^ algorithm and set the resolution based on each dataset’s number of cell types.

### Normalized mutual information (NMI)

NMI is also used to measure clustering accuracy and is defined as:

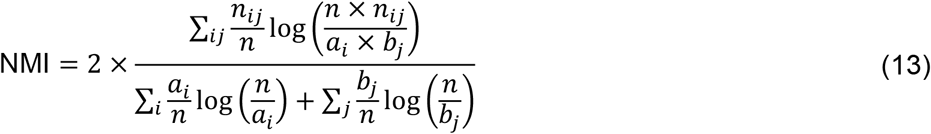

The notations are the same as those in ARI. NMI ranges in [0, 1] and higher values also indicate higher similarities between the clustering result and actual cell types. It can be calculated by the function ‘*normalized_mutual_info_score’* in the Python module *sklearn*.*metrics*.*cluster*. To calculate NMI, we utilized the *Louvain* algorithm and set the resolution based on the number of cell types in each dataset.

### Average silhouette width (ASW)

We employed the Average silhouette width (ASW) in conjunction with cell type and batch labels to assess batch correction. The silhouette score of a data point is computed by subtracting its average distance to other members in the same cluster from its average distance to all members of the neighboring clusters and then dividing by the larger of the two values.

The resulting score ranges from − 1 to 1, where a high score denotes that the data point fits well in the current cluster, while a low score indicates a poor fit. The average score of all data points is used to measure overall cell type purity or batch mixing through the choice of labels.

#### ASW_celltype

The silhouette width for cell type label (ASW_celltype) of cell *i* is defined as follows:

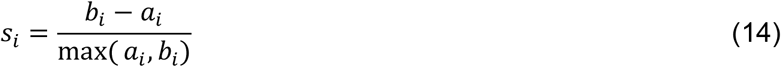

In this context, *a*_*i*_ represents the average distance from cell i to all other cells sharing the same label, while *b*_*i*_ denotes the minimum average distance from cell i to each group of cells assigned a different label. The ASW_celltype is calculated as the mean of the silhouette widths across all cells, where higher values indicate that cells are closer to cells with the same label and more distant from cells with different labels.

#### ASW_batch

The average silhouette width for batches (ASW_batch) is employed to assess the effectiveness of global batch mixing. The silhouette width for the batch of cell i is defined as follows:

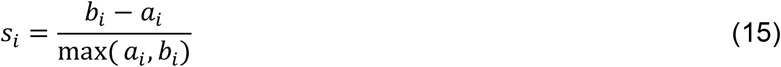

In this context, *a*_*i*_ represents the average distance from cell i to all other cells within the same batch, while *b*_*i*_ is the minimum average distance from cell i to groups of cells assigned to other batches. The ASW_batch is calculated as the mean of the silhouette widths across all cells, where higher values suggest that cells are more closely associated with others from the same batch and more distant from those in different batches. To ensure that higher scores reflect better batch mixing, the ASW_batch scores are adjusted by subtracting them from 1. Both metrics (ASW_celltype and ASW_batch) were computed on the embeddings provided by integration methods or the PCA of expression matrices in case of feature output. We utilized the ‘*silhouette_score’* function from the *sklearn*.*metrics* module in Python to calculate the ASW. Both ASW scores were then scaled to range between 0 and 1.

## Databases

### Tabula Muris transcriptomic mouse cell atlas

In this study, we utilize the Tabula Muris transcriptomic cell atlas^23^, which provides a comprehensive single-cell transcriptomic dataset comprising nearly 100000 cells across 20 mouse organs and tissues. To rigorously assess our annotation methodology, we applied SAFAARI to eight distinct organs, including the bladder, heart, kidney, liver, mammary gland, bone marrow, and muscle. In one scenario, for each organ, the 10x dataset served as the reference (source domain), while the FACS dataset was designated as the query (target domain). In a complementary analysis, the FACS dataset was employed as the reference, with the 10x dataset functioning as the target domain. This dual approach facilitates a robust evaluation of annotation accuracy across differing sequencing platforms. After preprocessing and normalization of the datasets, the intersection of highly variable genes of reference and target domains was selected for downstream analysis.

### Human pancreatic islet datasets

We conducted a multi-source integration study using several publicly available datasets on human pancreatic islets obtained from distinct studies and sequencing platforms including Fluidigm C1^30^ (GSE86469), CelSeq2^32^ (GSE85241), and Smart-Seq2^33^ (E-MTAB-5061). This approach enhances the richness of the reference domain, facilitating more robust label transfer. Following the preprocessing and normalization of these datasets, we identified 4,000 highly variable genes within each dataset and selected the intersection of these genes for subsequent analyses. The integrated dataset was then utilized as a reference to annotate the InDrops^31^ (GSE84133) dataset in an open-set scenario.

### Human Lung Atlas

We retrieved the single-cell expression data from the healthy human lung atlas, as detailed in the benchmarking atlas-level work^38^. This atlas encompasses samples from three different laboratories, generated using both Drop-seq and 10x Chromium platforms. The Drop-seq data was sourced from GEO under accession code GSE130148. We considered the transplant datasets as the reference domain, which includes 10x along with Drop-seq data and the 10x biopsy dataset as the target domain. It should be mentioned that lung biopsy data comes from a distinct spatial location (the airways) relative to the location of transplant samples (the parenchyma)^38^. We selected the intersection of highly variable genes from the transplant and biopsy datasets for open-set label transfer^38^. Following preprocessing and normalization, we selected the intersection of highly variable genes from the transplant and biopsy datasets for subsequent analysis of open-set label transfer.

### Human ovary multi-omics dataset

To evaluate SAFAARI’s capability in integrating ATAC-seq and RNA-seq data, we utilized datasets from a study^48^, which provides an epigenomic perspective, enabling the study of chromatin accessibility alongside gene expression in ovarian cells and includes transcriptomic (scRNA-seq) and regulatory (scATAC-seq) profiles of the normal postmenopausal ovary and fallopian tube to understand the cellular composition and gene regulatory networks in these tissues, which are crucial for understanding gynecological diseases in menopause, such as ovarian cancer. Following preprocessing and normalization, two datasets were prepared for downstream analysis: an ATAC-seq dataset comprising 18315 cells and an RNA-seq dataset with 26087 cells.

### Human PBMC multi-omics dataset

We utilized a single cell multiome dataset from peripheral blood mononuclear cells (PBMCs) obtained from a cryopreserved sample of a healthy female donor by 10x Genomics from AllCells company^49^. This dataset offers paired ATAC-seq and gene expression profiles, enabling a comprehensive analysis of chromatin accessibility alongside transcriptomic data. This experiment further evaluated SAFAARI’s capability to integrate ATAC-seq and RNA-seq data effectively.

For preprocessing and normalization of the ATAC-seq data, we used the Signac package and followed established pipelines for ATAC-seq integration, as described in Seurat v5. After these steps, two batches of data were prepared, comprising approximately 11000 cells each for ATAC-seq and RNA-seq. (https://satijalab.org/seurat/articles/seurat5_atacseq_integration_vignette)

### Simulated datasets

For the simulation tasks, two datasets were generated using the Splatter package^40^ were retrieved from a previous benchmarking study^38^ to assess data integration methods in a controlled setting.

#### Simulated dataset 1

This dataset comprises six batches designed to replicate an experiment involving multiple samples derived from a single tissue containing seven cell types, each generated using distinct technologies. This simulation poses several challenges for data integration, as the batches vary in the number of cells (ranging from 1000 to 3000), cell type proportions (0–35%), and counts per cell (30–100% of the baseline)^38^.

#### Simulated dataset 2

This dataset is designed to mimic a more complex experimental setup with a nested structure comprising four batches, each containing three sub-batches. This design reflects a multi-center study, where each center processes multiple batches, potentially using different technologies. In this scenario, the batch effect between centers is expected to be more pronounced than the batch effect between sub-batches within the same center. This additional layer of variation introduces a challenge for integration methods, which must account for both batch and sub-batch effects while preserving distinctions between cell types. Furthermore, the limited number of cells available to estimate batch effects adds an additional complexity to this task. The number of groups in Simulation 2 has been reduced to four^38^.

### Human PBMC CITE-seq reference atlas

This CITE-seq reference comprises a comprehensive dataset of 162000 human peripheral blood mononuclear cells (PBMCs), measured with 228 antibodies. This multimodal dataset was constructed using a technique called Cellular Indexing of Transcriptomes and Epitopes by Sequencing (CITE-seq), which combines RNA sequencing with simultaneous protein measurements through antibody labeling. This approach enhances the ability to capture both transcriptomic and proteomic data, enabling more detailed identification of cell states. The weighted-nearest neighbor analysis applied to this dataset improves the resolution of cellular subpopulations, allowing for the identification of previously unreported lymphoid subtypes. It serves as a robust multimodal reference atlas, offering a more unified understanding of cellular identity through the integration of multiple data modalities^34^. Following preprocessing and normalization, two batches (protein and RNA) were prepared for integration.

### Human kidney scATAC-seq datasets

To evaluate SAFAARI’s capability for annotating single-nucleus ATAC-seq data in an open-set scenario, we utilized two snATAC-seq datasets from studies of the kidney cortex: the Muto *et al*. study^44^ and the Wilson *et al*. study^45^. Control samples were extracted from both studies, with the Muto dataset comprising samples from patients undergoing nephrectomy (n=5) while the Wilson dataset also includes a sample from a deceased organ donor (n=6). After preprocessing and normalization, the Muto *et al*. dataset, comprising 24835 cells, was designated as the reference domain, while the Wilson *et al*. dataset, containing 36732 cells, was used as the target domain. SAFFARI was then applied to assess its performance in this open-set scenario.

### Cross-species muscle datasets

To assess the robustness of SAFAARI in retaining species-specific features while identifying conserved cell types across species, we utilized three single-cell RNA sequencing (scRNA-seq) datasets from studies on muscle tissue in mice, bovine, and humans. The mouse dataset, from Dell’Orso et al.^41^ examined skeletal muscle stem cells (MuSCs) in adult mice under homeostatic and regenerative conditions, offering insights into MuSC contributions to muscle repair following injury. For bovine data, we used Messmer et al.^42^ who performed scRNA-seq on bovine muscle-derived cells to advance cultured meat production by profiling cellular heterogeneity and optimizing growth conditions. Lastly, the human dataset from Rubenstein et al.^43^ provided a detailed transcriptional map of human skeletal muscle, identifying key cell types and highlighting tissue heterogeneity. We configured the open-set annotation settings to map cell types from mice to humans and bovine to humans.

## Data and Code Availability

All datasets can be accessed through the respective links that are listed in SAFAARI GitHub repository: https://github.com/VafaeeLab/SAFAARI/blob/main/DataLinks.md. Details of these datasets and their usage are available in the Supplementary Tables. An open-source implementation of the SAFAARI algorithm is available at https://github.com/VafaeeLab/SAFAARI.

## Supplementary Tables

**Supplementary Table 1**. Summary of Tabula Muris Transcriptomic Mouse Cell Atlas (Page 2)

**Supplementary Table 2**. Summary of Human Pancreatic Islet Datasets (Page 4)

**Supplementary Table 3**. Summary of Simulated Datasets (Page 5)

**Supplementary Table 4**. Summary of Human Lung Atlas (Page 5)

**Supplementary Table 5**. Summary of Cross-species Muscle Datasets (Page 6)

**Supplementary Table 6**. Summary of Human Kidney scATAC seq Datasets (Page 6)

**Supplementary Table 7**. Summary of Human Ovary Multi-Omics Dataset (Page 7)

**Supplementary Table 8**. Summary of Human PBMC Multi-Omics Dataset (Page 7)

**Supplementary Table 9**. Summary of Human PBMC CITE-seq Reference Atlas (Page7)

**Supplementary Table 1.** The scRNA-seq data from eight different tissues in the Tabula Muris cell atlas [1] (GSE132042) was obtained where the gene counts were derived using two techniques: 10X Genomics (10X) and FACS-based cell capture in plates (FACS). The table lists the cell types and the number of cells from both 10X and FACS datasets under two scenarios: (1) 10X is considered as the reference (or source) dataset and FACS as the query (or target) dataset, and (2) the reverse, with FACS as the reference and 10X as target (c.f., ‘Domain’ column).

| Organ | Domain | Platform | Cell Types | No. of Cells | Cell type (No. of Cells) |
| --- | --- | --- | --- | --- | --- |
| Bladder | Reference | 10X | 2 | 2109 | bladder cell (923), mesenchymal cell (1186), basal cell of urothelium (0) |
|  | Target | FACS | 3 | 1287 | bladder cell (532), mesenchymal cell (656), basal cell of urothelium: *unknown* (99) |
|  | Reference | FACS | 2 | 1188 | bladder cell (532), mesenchymal cell (656), leukocyte(0) |
|  | Target | 10X | 3 | 2109 | bladder cell (923), mesenchymal cell (1186), leukocyte: *unknown* (57) |
| Heart | Reference | 10X | 6 | 624 | cardiac muscle cell (83), endocardial cell (65), endothelial cell (178), erythrocyte (55), fibroblast (222), smooth muscle cell (21), epicardial adipocyte (0) |
|  | Target | FACS | 7 | 4029 | cardiac muscle cell (140), endocardial cell (175), endothelial cell (1268), erythrocyte (11), fibroblast (2087), smooth muscle cell (255), epicardial adipocyte: *unknown* (93) |
|  | Reference | FACS | 5 | 3681 | cardiac muscle cell (140), endocardial cell (175), endothelial cell (1268), erythrocyte (11), fibroblast (2087), smooth muscle cell (0) |
|  | Target | 10X | 6 | 624 | cardiac muscle cell (83), endocardial cell (65), endothelial cell (178), erythrocyte (55), fibroblast (222), smooth muscle cell: *unknown* (21) |
| Kidney | Reference | 10X | 5 | 2496 | endothelial cell (45), fenestrated cell (345), fibroblast (113), kidney collecting duct cell (72), kidney tubule cell (1921), leukocyte (0) |
|  | Target | FACS | 6 | 517 | endothelial cell (40), fenestrated cell (69), fibroblast (65), kidney collecting duct cell (44), kidney tubule cell (261), leukocyte: *unknown* (38) |
|  | Reference | FACS | 5 | 479 | endothelial cell (40), fenestrated cell (69), fibroblast (65), kidney collecting duct cell (44), kidney tubule cell (261), leukocyte (0) |
|  | Target | 10X | 6 | 2550 | endothelial cell (45), fenestrated cell (345), fibroblast (113), kidney collecting duct cell (72), kidney tubule cell (1921), leukocyte: *unknown* (54) |
| Liver | Reference | 10X | 2 | 1026 | endothelial cell (20), hepatocyte (1006), B cell (0) |
|  | Target | FACS | 3 | 624 | endothelial cell (196), hepatocyte (399), B cell: *unknown* (29) |
| Mammary | Reference | 10X | 3 | 1341 | basal cell (393), endothelial cell (251), stromal cell (697), luminal epithelial cell of mammary gland (0) |
|  | Target | FACS | 4 | 2304 | basal cell (1275), endothelial cell (49), stromal cell (428), luminal epithelial cell of mammary gland: *unknown* (552) |
|  | Reference | FACS | 3 | 1752 | basal cell (1275), endothelial cell (49), stromal cell (428), macrophage (0) |
|  | Target | 10X | 4 | 1527 | basal cell (393), endothelial cell (251), stromal cell (697), macrophage: *unknown* (186) |
| Marrow | Reference | 10X | 6 | 3288 | B cell (390), Fraction A pre-pro B cell (63), T cell (162), granulocyte (770), hematopoietic stem cell (1381), monocyte (522), natural killer cell (0) |
|  | Target | FACS | 7 | 4298 | B cell (1848), Fraction A pre-pro B cell (177), T cell (142), granulocyte (380), hematopoietic stem cell (1291), monocyte (324), natural killer cell: *unknown* (136) |
|  | Reference | FACS | 6 | 4162 | B cell (1848), Fraction A pre-pro B cell (177), T cell (142), granulocyte (380), hematopoietic stem cell (1291), monocyte (324), erythrocyte (0) |
|  | Target | 10X | 7 | 3430 | B cell (390), Fraction A pre-pro B cell (63), T cell (162), granulocyte (770), hematopoietic stem cell (1381), monocyte (522), erythrocyte: *unknown* (142) |
| Muscle | Reference | 10X | 6 | 3844 | B cell (463), T cell (323), endothelial cell (1290), macrophage (308), mesenchymal stem cell (1111), skeletal muscle satellite cell (349), skeletal muscle satellite stem cell (0) |
|  | Target | FACS | 7 | 1937 | B cell (151), T cell (35), endothelial cell (206), macrophage (71), mesenchymal stem cell (486), skeletal muscle satellite cell (546), skeletal muscle satellite stem cell: *unknown* (442) |
|  | Reference | FACS | 6 | 1495 | B cell (151), T cell (35), endothelial cell (206), macrophage (71), mesenchymal stem cell (486), skeletal muscle satellite cell (546), chondroblast (0) |
|  | Target | 10X | 7 | 4224 | B cell (463), T cell (323), endothelial cell (1290), macrophage (308), mesenchymal stem cell (1111), skeletal muscle satellite cell (349), chondroblast: *unknown* (380) |
| Spleen | Reference | 10X | 2 | 9013 | B cell (6726), T cell (2287), myeloid cell (0) |
|  | Target | FACS | 3 | 1689 | B cell (1238), T cell (403), myeloid cell: *unknown* (48) |
|  | Reference | FACS | 2 | 1641 | B cell (1238), T cell (403), dendritic cell (0) |
|  | Target | 10X | 3 | 9157 | B cell (6726), T cell (2287), dendritic cell: *unknown* (144) |

**Supplementary Table 2.** We obtained human pancreas datasets from distinct studies and platforms to apply open set annotation. We consider Lawlor, Muraro and Segerstolpe as the source domain and annotated Baron dataset as the target domain.

| Organ | Domain | 1<sup>st</sup> Author [Ref] | Platform | Cell types | No. of Cells | Accession | Cell type (No. of Cells) |
| --- | --- | --- | --- | --- | --- | --- | --- |
| Pancreas | Reference | Lawlor [2] | Fluidigm C1 | 7 | 617 | GSE86469 | Betta (264), Ductal (28), Alpha (239), Acinar (24), Gamma (18), Delta (25), Stellate (19), Macrophage: *unknown* (0) |
|  | Reference | Muraro [3] | CelSeq2 | 9 | 2106 | GSE85241 | Betta (446), Ductal (244), Alpha (808), Acinar (218), Gamma (97), Delta (191), Endothelial (21), Epsilon (3), Mesenchymal (78), Macrophage: *unknown* (0) |
|  | Reference | Segerstolpe [4] | Smart-Seq2 | 10 | 2122 | E-MTAB-5061 | Betta (270), Ductal (386), Alpha (886), Acinar (185), Gamma (197), Delta (114), Endothelial (16), Epsilon (7), Mast (7), PSC (54), Macrophage: *unknown* (0) |
|  | Target | Baron [5] | InDrop | 8 | 8254 | GSE84133 | Betta (2525), Ductal (1077), Alpha (2326), Acinar (958), Gamma (255), Delta (601), Stellate (457), Macrophage: *unknown* (55) |

**Supplementary Table 3.** There are two simulated datasets. Simulated dataset1 comprises six batches, designed to replicate an experiment involving multiple samples derived from a single tissue containing seven cell types, each generated using distinct technologies. Simulated dataset 2 is designed to mimic a more complex experimental setup with a nested structure comprising four batches, each containing three sub-batches. This design reflects a multi-center study, where each center processes multiple batches, potentially using different technologies. We integrated different batches in both simulated datasets.

| Dataset Name [Ref] | Batch | Cell Types (Groups) | No. of Cells | Cell Types (No. of Cells) |
| --- | --- | --- | --- | --- |
| Simulated Dataset 1 [6] | Batch 1 | 6 | 2908 | Group 1 (1026), Group 2 (639), Group 3 (440), Group 4 (372), Group 5 (288), Group 6 (143) |
|  | Batch 2 | 6 | 2422 | Group 1 (601), Group 2 (803), Group 3 (487), Group 4 (240), Group 6 (194), Group 7 (97) |
|  | Batch 3 | 5 | 2120 | Group 1 (634), Group 2 (638), Group 4 (425), Group 5 (214), Group 6 (209) |
|  | Batch 4 | 5 | 1929 | Group 1 (288), Group 2 (387), Group 3 (681), Group 4 (285), Group 5 (288) |
|  | Batch 6 | 4 | 1761 | Group 1 (433), Group 2 (331), Group 3 (144), Group 4 (49) |
|  | Batch 5 | 7 | 957 | Group 1 (706), Group 2 (87), Group 3 (177), Group 4 (175), Group 5 (299), Group 6 (210), Group 7(107) |
| Simulated Dataset 2 [6] | Batch 1 | 3 | 4806 | Group 1 (1682), Group 2 (1058), Group 3 (1212), Group 4 (854) |
|  | Batch 2 | 3 | 4905 | Group 1 (1198), Group 2 (1921), Group 3 (1303), Group 4 (483) |
|  | Batch 3 | 3 | 4809 | Group 1 (1446), Group 2 (726), Group 3 (1673), Group 4: (964) |
|  | Batch 4 | 4 | 4798 | Group 1 (481), Group 2 (1447), Group 3 (953), Group 4 (1917) |

**Supplementary Table 4.**
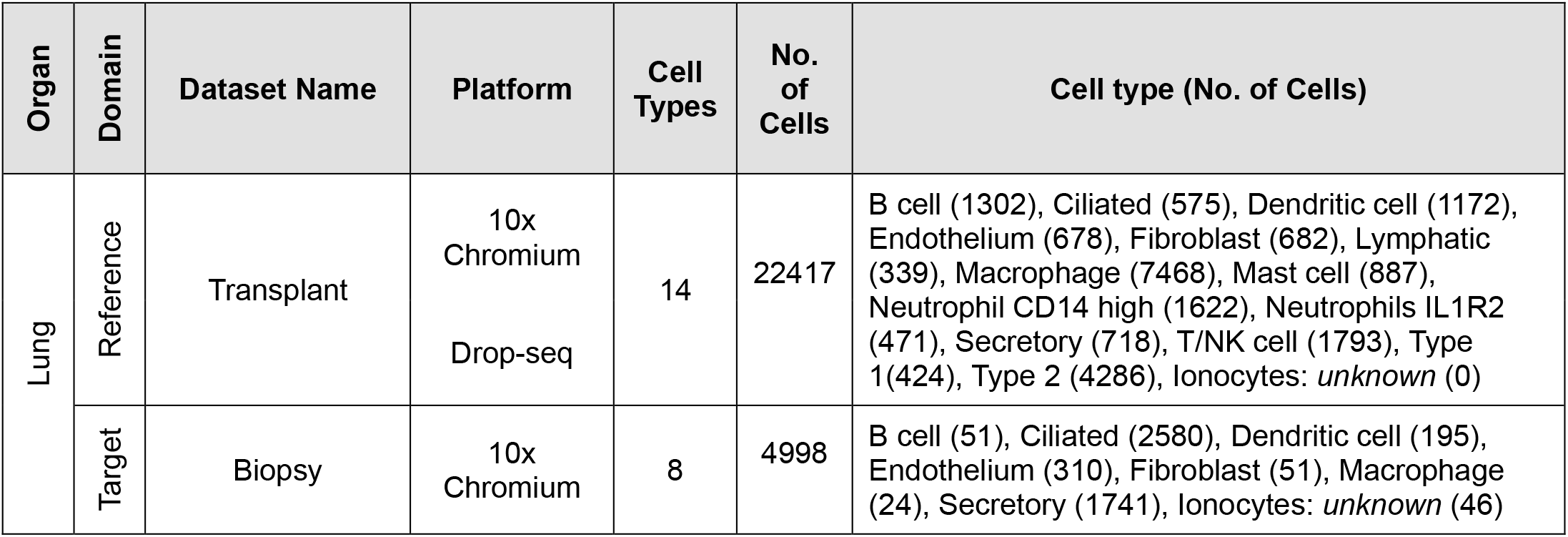
We utilized the single-cell expression data from the human lung atlas, as detailed in the work of [6]. This atlas encompasses samples from three different laboratories, generated using both Drop-seq and 10X Chromium platforms. The Drop-seq data was sourced from GEO under accession code GSE130148. For our experiment, we considered the transplant datasets as the reference domain and the biopsy dataset as the target domain.

**Supplementary Table 5.**
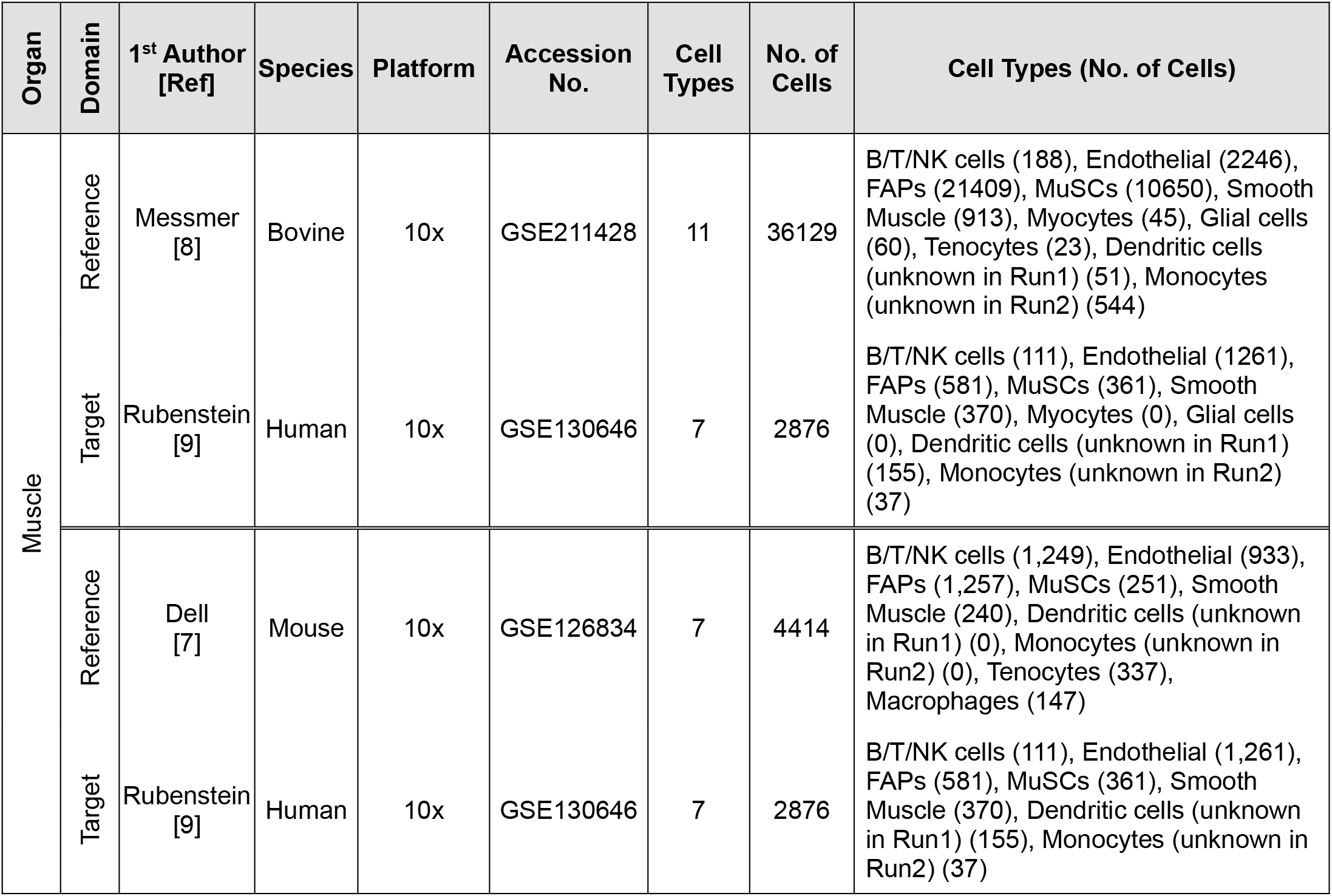
We used three datasets from different studies on muscle tissue from mouse[7], bovine [8], and human [9]. We configured the open set annotation settings to map from mouse to human and from bovine to human, running two separate experiments where different cell types were treated as unknown.

| Organ | Domain | 1<sup>st</sup> Author [Ref] | Species | Platform | Accession No. | Cell Types | No. of Cells | Cell Types (No. of Cells) |
| --- | --- | --- | --- | --- | --- | --- | --- | --- |
| Muscle | Reference | Messmer [8] | Bovine | 10x | GSE211428 | 11 | 36129 | B/T/NK cells (188), Endothelial (2246), FAPs (21409), MuSCs (10650), Smooth Muscle (913), Myocytes (45), Glial cells (60), Tenocytes (23), Dendritic cells (unknown in Run1) (51), Monocytes (unknown in Run2) (544) |
|  | Target | Rubenstein [9] | Human | 10x | GSE130646 | 7 | 2876 | B/T/NK cells (111), Endothelial (1261), FAPs (581), MuSCs (361), Smooth Muscle (370), Myocytes (0), Glial cells (0), Dendritic cells (unknown in Run1) (155), Monocytes (unknown in Run2) (37) |
|  | Reference | Dell [7] | Mouse | 10x | GSE126834 | 7 | 4414 | B/T/NK cells (1,249), Endothelial (933), FAPs (1,257), MuSCs (251), Smooth Muscle (240), Dendritic cells (unknown in Run1) (0), Monocytes (unknown in Run2) (0), Tenocytes (337), Macrophages (147) |
|  | Target | Rubenstein [9] | Human | 10x | GSE130646 | 7 | 2876 | B/T/NK cells (111), Endothelial (1,261), FAPs (581), MuSCs (361), Smooth Muscle (370), Dendritic cells (unknown in Run1) (155), Monocytes (unknown in Run2) (37) |

**Supplementary Table 6.** We utilize two sc-ATAC-seq datasets from studies on the kidney cortex: the Muto dataset [10] and the Wilson dataset [11]. We annotated Wilson as the target dataset according to Muto Dataset which was the source dataset.

| Organ | Domain | 1<sup>st</sup> Author [Ref] | Sequencing Method | Platform | Cell Types | No. of Cells | Cell type (No. of Cells) |
| --- | --- | --- | --- | --- | --- | --- | --- |
| Kidney | Reference | Muto [10] | ATAC-seq | 10X | 10 | 24835 | Podocyte (133), Renal beta-intercalated cell (620), Epithelial cell of proximal tubule (690), Renal principal cell (1,291), Renal alpha-intercalated cell (613), Parietal epithelial cell (402), Kidney proximal convoluted tubule epithelial cell (6384), Kidney proximal straight tubule epithelial cell (4146), Kidney distal convoluted tubule epithelial cell (2781), Kidney loop of Henle thick ascending limb epithelial cell (7775), Endothelial cell: unknown (0) |
|  | Target | Wilson [11] | ATAC-seq | 10X | 11 | 36732 | Podocyte (241), Renal beta-intercalated cell (729), Epithelial cell of proximal tubule (1,734), Renal principal cell (975), Renal alpha-intercalated cell (1112), Parietal epithelial cell (578), Kidney proximal convoluted tubule epithelial cell (12467), Kidney proximal straight tubule epithelial cell (3451), Kidney distal convoluted tubule epithelial cell (5995), Kidney loop of Henle thick ascending limb epithelial cell (8176), Endothelial cell: unknown (1274) |

**Supplementary Table 7.** We utilized datasets from the study by [12], which dataset provides an epigenomic perspective, enabling the study of chromatin accessibility alongside gene expression in human ovarian cells and includes transcriptomic (scRNA-seq) and regulatory (scATAC-seq) profiles of the normal postmenopausal ovary and fallopian tube. We focused on the ovary and integrated RNA-seq and ATAC-seq datasets.

| Organ | Dataset Name | Platform | Cell Types | No. of Cells | Cell type (No. of Cells) |
| --- | --- | --- | --- | --- | --- |
| Ovary | RNA-seq | Drop-seq | 5 | 26087 | Endothelial cell (291), Smooth muscle cell (150), Stromal cell (21572), Pericyte (3483), Leukocyte (591) |
|  | ATAC-seq | scATAC-seq | 5 | 18315 | Endothelial cell (156), Smooth muscle cell (165), Stromal cell (15607), Pericyte (2,104), Leukocyte (283) |

**Supplementary Table 8.**
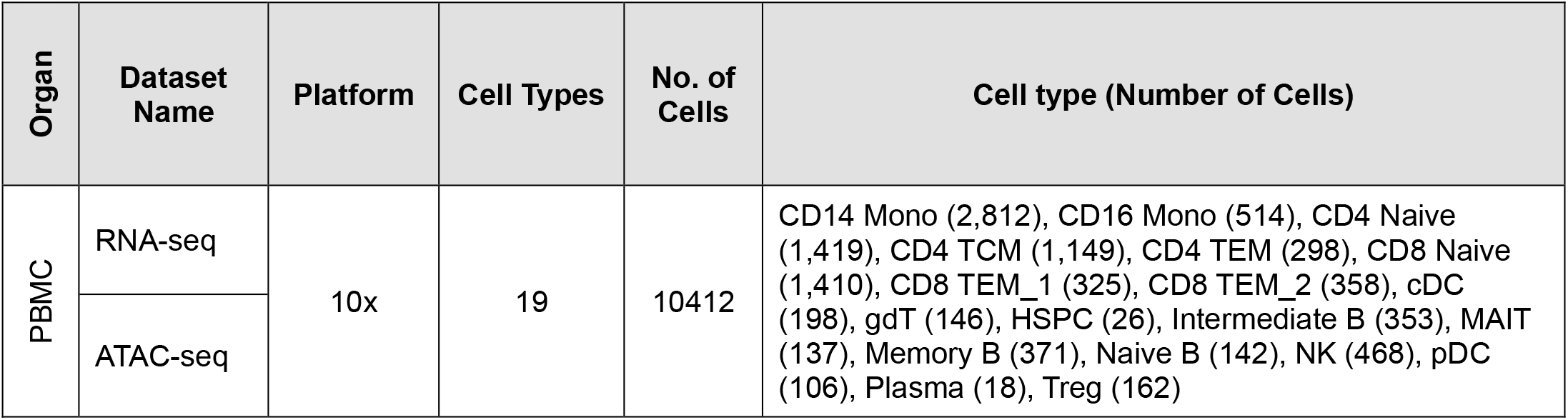
We utilized a single cell multiome dataset from peripheral blood mononuclear cells (PBMCs) [13] obtained from a cryopreserved sample of a healthy female donor. This dataset offers paired ATAC-seq and gene expression profiles, enabling a comprehensive analysis of chromatin accessibility alongside transcriptomic data. We integrated RNA-seq and ATAC-seq datasets.

**Supplementary Table 9.** This CITE-seq reference consists of a comprehensive dataset of 162000 human peripheral blood mononuclear cells (PBMCs), measured with 228 antibodies [14]. This multimodal dataset was constructed using a technique called Cellular Indexing of Transcriptomes and Epitopes by Sequencing (CITE-seq), which combines RNA sequencing with simultaneous protein measurements through antibody labeling, which were our datasets to be integrated.

| Organ | Dataset Name | Platform | Cell Types | No. of Cells | Cell type (Number of Cells) |
| --- | --- | --- | --- | --- | --- |
| PBMC | Protein | CITE-seq | 8 | 161764 | B (13800), CD4 T (41001), CD8 T (25469), DC (3589), Mono (49010), NK (18664), Other (3442), Other T (6789) |
|  | RNA |  |  |  |  |

## Notes

### Competing Interest Statement

The authors have declared no competing interest.

## References

1 Lopez, R., Regier, J., Cole, M. B., Jordan, M. I. & Yosef, N. Deep generative modeling for single-cell transcriptomics. Nature Methods 15, 1053–1058 (2018). 10.1038/s41592-018-0229-2

2 Yang, M. et al. Contrastive learning enables rapid mapping to multimodal single-cell atlas of multimillion scale. Nature Machine Intelligence 4, 696–709 (2022). 10.1038/s42256-022-00518-z

3 Chen, H., Ryu, J., Vinyard, M. E., Lerer, A. & Pinello, L. SIMBA: single-cell embedding along with features. Nature Methods 21, 1003–1013 (2024). 10.1038/s41592-023-01899-8

4 Brbić, M. et al. Annotation of spatially resolved single-cell data with STELLAR. Nature Methods 19, 1411–1418 (2022). 10.1038/s41592-022-01651-8

5 Shao, X. et al. scDeepSort: a pre-trained cell-type annotation method for single-cell transcriptomics using deep learning with a weighted graph neural network. Nucleic Acids Research 49, e122–e122 (2021). 10.1093/nar/gkab775

6 Fischer, F. et al. scTab: Scaling cross-tissue single-cell annotation models. Nature Communications 15, 6611 (2024). 10.1038/s41467-024-51059-5

7 Lotfollahi, M. et al. Mapping single-cell data to reference atlases by transfer learning. Nature Biotechnology, 1–10 (2021).

8 Rosen, Y. et al. Toward universal cell embeddings: integrating single-cell RNA-seq datasets across species with SATURN. Nature Methods 21, 1492–1500 (2024). 10.1038/s41592-024-02191-z

9 Yang, X., Mann Koren K., Wu, H. & Ding, J. scCross: a deep generative model for unifying single-cell multi-omics with seamless integration, cross-modal generation, and in silico exploration. Genome Biology 25, 198 (2024). 10.1186/s13059-024-03338-z

10 Ren, P. et al. Single-cell assignment using multiple-adversarial domain adaptation network with large-scale references. Cell Rep Methods 3, 100577 (2023). 10.1016/j.crmeth.2023.100577

11 Kiselev, V. Y., Andrews, T. S. & Hemberg, M. Challenges in unsupervised clustering of single-cell RNA-seq data. Nature Reviews Genetics 20, 273–282 (2019). 10.1038/s41576-018-0088-9

12 Yang, F. et al. scBERT as a large-scale pretrained deep language model for cell type annotation of single-cell RNA-seq data. Nature Machine Intelligence 4, 852–866 (2022).

13 Hou, W. & Ji, Z. J. N. M. Assessing GPT-4 for cell type annotation in single-cell RNA-seq analysis. Nature Methods, 1–4 (2024).

14 Zhang, Y. et al. CellSTAR: a comprehensive resource for single-cell transcriptomic annotation. Nucleic Acids Research 52, D859–D870 (2023). 10.1093/nar/gkad874

15 Hu, J. et al. Iterative transfer learning with neural network for clustering and cell type classification in single-cell RNA-seq analysis. Nature Machine Intelligence 2, 607–618 (2020). 10.1038/s42256-020-00233-7

16 Farahani, A., Voghoei, S., Rasheed, K. & Arabnia, H. R. A brief review of domain adaptation. Advances in data science information engineering, 877–894 (2021).

17 Wu, J. & He, J. Domain Adaptation with Dynamic Open-Set Targets, in Proceedings of the 28th ACM SIGKDD Conference on Knowledge Discovery and Data Mining. 2039–2049 (ACM).

18 Fernández, A., Garcia, S., Herrera, F. & Chawla, N. V. J. J. o. a. i. r. SMOTE for learning from imbalanced data: progress and challenges, marking the 15-year anniversary. Journal of artificial intelligence research 61, 863–905 (2018).

19 Zheng, L., Xiong, J., Zhu, Y. & He, J. Contrastive Learning with Complex Heterogeneity, in Proceedings of the 28th ACM SIGKDD Conference on Knowledge Discovery and Data Mining. 2594–2604.

20 Khosla, P. et al. Supervised contrastive learning. Advances in neural information processing systems 33, 18661–18673 (2020).

21 Gunawan, I., Vafaee, F., Meijering, E. & Lock, J. G. J. C. R. M. An introduction to representation learning for single-cell data analysis. Cell Reports Methods 3 (2023).

22 Qu, H.-Q., Kao, C. & Hakonarson, H. Single-Cell RNA Sequencing Technology Landscape in 2023. Stem Cells 42, 1–12 (2023). 10.1093/stmcls/sxad077

23 Schaum, N. et al. Single-cell transcriptomics of 20 mouse organs creates a Tabula Muris: The Tabula Muris Consortium. Nature 562, 367 (2018).

24 Abdelaal, T. et al. A comparison of automatic cell identification methods for single-cell RNA sequencing data. Genome biology 20, 1–19 (2019).

25 Breiman, L. Random forests. Machine learning 45, 5–32 (2001).

26 Ma, W., Su, K. & Wu, H. Evaluation of some aspects in supervised cell type identification for single-cell RNA-seq: classifier, feature selection, and reference construction. Genome Biology 22, 264 (2021). 10.1186/s13059-021-02480-2

27 Chen, X. et al. Cell type annotation of single-cell chromatin accessibility data via supervised Bayesian embedding. Nature Machine Intelligence 4, 116–126 (2022). 10.1038/s42256-021-00432-w

28 Stuart, T. et al. Comprehensive Integration of Single-Cell Data. Cell 177, 1888–1902.e1821 (2019). 10.1016/j.cell.2019.05.031

29 Aran, D. et al. Reference-based analysis of lung single-cell sequencing reveals a transitional profibrotic macrophage. Nature Immunology 20, 163–172 (2019). 10.1038/s41590-018-0276-y

30 Lawlor, N. et al. Single-cell transcriptomes identify human islet cell signatures and reveal cell-type-specific expression changes in type 2 diabetes. Genome Res 27, 208–222 (2017). 10.1101/gr.212720.116

31 Baron, M. et al. A Single-Cell Transcriptomic Map of the Human and Mouse Pancreas Reveals Inter- and Intra-cell Population Structure. Cell Syst 3, 346–360.e344 (2016). 10.1016/j.cels.2016.08.011

32 Muraro, M. J. et al. A Single-Cell Transcriptome Atlas of the Human Pancreas. Cell Syst 3, 385–394.e383 (2016). 10.1016/j.cels.2016.09.002

33 Segerstolpe, Å. et al. Single-Cell Transcriptome Profiling of Human Pancreatic Islets in Health and Type 2 Diabetes. Cell Metab 24, 593–607 (2016). 10.1016/j.cmet.2016.08.020

34 Hao, Y. et al. Integrated analysis of multimodal single-cell data. Cell 184, 3573–3587.e3529 (2021). 10.1016/j.cell.2021.04.048

35 Korsunsky, I. et al. Fast, sensitive and accurate integration of single-cell data with Harmony. Nature methods 16, 1289–1296 (2019).

36 Hie, B., Bryson, B. & Berger, B. Efficient integration of heterogeneous single-cell transcriptomes using Scanorama. Nature biotechnology 37, 685–691 (2019).

37 Polański, K. et al. BBKNN: fast batch alignment of single cell transcriptomes. Bioinformatics 36, 964–965 (2020).

38 Luecken, M. D. et al. Benchmarking atlas-level data integration in single-cell genomics. Nature Methods 19, 41–50 (2022). 10.1038/s41592-021-01336-8

39 Tran, H. T. N. et al. A benchmark of batch-effect correction methods for single-cell RNA sequencing data. Genome biology 21, 1–32 (2020).

40 Zappia, L., Phipson, B. & Oshlack, A. Splatter: simulation of single-cell RNA sequencing data. Genome Biology 18, 174 (2017). 10.1186/s13059-017-1305-0

41 Dell’Orso, S. et al. Single cell analysis of adult mouse skeletal muscle stem cells in homeostatic and regenerative conditions. Development 146 (2019). 10.1242/dev.174177

42 Messmer, T. et al. Single-cell analysis of bovine muscle-derived cell types for cultured meat production. Frontiers in nutrition 10, 1212196 (2023).

43 Rubenstein, A. B. et al. Single-cell transcriptional profiles in human skeletal muscle. Scientific Reports 10, 229 (2020). 10.1038/s41598-019-57110-6

44 Muto, Y. et al. Single cell transcriptional and chromatin accessibility profiling redefine cellular heterogeneity in the adult human kidney. Nature Communications 12, 2190 (2021). 10.1038/s41467-021-22368-w

45 Wilson, P. C. et al. Multimodal single cell sequencing implicates chromatin accessibility and genetic background in diabetic kidney disease progression. Nature Communications 13, 5253 (2022). 10.1038/s41467-022-32972-z

46 Xiao, C., Chen, Y., Meng, Q., Wei, L. & Zhang, X. Benchmarking multi-omics integration algorithms across single-cell RNA and ATAC data. Briefings in Bioinformatics 25 (2024). 10.1093/bib/bbae095

47 Baysoy, A., Bai, Z., Satija, R. & Fan, R. The technological landscape and applications of single-cell multi-omics. Nature Reviews Molecular Cell Biology 24, 695–713 (2023). 10.1038/s41580-023-00615-w

48 Lengyel, E. et al. A molecular atlas of the human postmenopausal fallopian tube and ovary from single-cell RNA and ATAC sequencing. Cell Rep 41, 111838 (2022). 10.1016/j.celrep.2022.111838

49 Genomics, x. (ed 10x Genomics) (10x Genomics, 2020).

50 Cao, Z.-J. & Gao, G. Multi-omics single-cell data integration and regulatory inference with graph-linked embedding. Nature Biotechnology 40, 1458–1466 (2022). 10.1038/s41587-022-01284-4

51 Jingyu, Z., Chen, Z., Wei, P., Li, G. & Lin, L. Open Set Domain Adaptation By Novel Class Discovery. (2022).

52 Pei, Z., Cao, Z., Long, M. & Wang, J. Multi-Adversarial Domain Adaptation. Proceedings of the AAAI Conference on Artificial Intelligence 32 (2018). 10.1609/aaai.v32i1.11767

53 Kiryo, R., Niu, G., Du Plessis, M. C. & Sugiyama, M. Positive-unlabeled learning with non-negative risk estimator. Advances in neural information processing systems 30 (2017).

54 Xu, Y., Xu, C., Xu, C. & Tao, D. in IJCAI. 3182–3188.

55 Ganin, Y. et al. Domain-adversarial training of neural networks. Journal of machine learning research 17, 1–35 (2016).

56 Robinson, M. D. & Oshlack, A. A scaling normalization method for differential expression analysis of RNA-seq data. Genome Biology 11, R25 (2010). 10.1186/gb-2010-11-3-r25

57 Van der Maaten, L. & Hinton, G. J. J. o. m. l. r. Visualizing data using t-SNE. Journal of machine learning research 9 (2008).

58 McInnes, L., Healy, J. & Melville, J. Umap: Uniform manifold approximation and projection for dimension reduction. arXiv preprint arXiv:1802.03426 (2018).

59 Rousseeuw, P. J. Silhouettes: a graphical aid to the interpretation and validation of cluster analysis. Journal of computational and applied mathematics 20, 53–65 (1987).

60 Hubert, L. & Arabie, P. Comparing partitions. Journal of classification 2, 193–218 (1985).

61 Danon, L., Diaz-Guilera, A., Duch, J. & Arenas, A. Comparing community structure identification. Journal of statistical mechanics: Theory and experiment 2005, P09008 (2005).

62 Blondel, V. D., Guillaume, J.-L., Lambiotte, R. & Lefebvre, E. Fast unfolding of communities in large networks. Journal of statistical mechanics: theory and experiment 2008, P10008 (2008).

## References

1. Schaum, N., et al., Single-cell transcriptomics of 20 mouse organs creates a Tabula Muris: The Tabula Muris Consortium. Nature, 2018. 562(7727): p. 367.

2. Lawlor, N., et al., Single-cell transcriptomes identify human islet cell signatures and reveal cell-type-specific expression changes in type 2 diabetes. Genome Res, 2017. 27(2): p. 208–222.

3. Muraro, M.J., et al., A Single-Cell Transcriptome Atlas of the Human Pancreas. Cell Syst, 2016. 3(4): p. 385–394.e3.

4. Segerstolpe, Å., et al., Single-Cell Transcriptome Profiling of Human Pancreatic Islets in Health and Type 2 Diabetes. Cell Metab, 2016. 24(4): p. 593–607.

5. Baron, M., et al., A Single-Cell Transcriptomic Map of the Human and Mouse Pancreas Reveals Inter- and Intra-cell Population Structure. Cell Syst, 2016. 3(4): p. 346–360.e4.

6 Luecken, M.D., et al., Benchmarking atlas-level data integration in single-cell genomics. Nat Methods, 2022. 19(1): p. 41–50.

7. Dell’Orso, S., et al., Single cell analysis of adult mouse skeletal muscle stem cells in homeostatic and regenerative conditions. Development, 2019. 146(12).

8. Messmer, T., et al., Single-cell analysis of bovine muscle-derived cell types for cultured meat production. Frontiers in nutrition, 2023. 10: p. 1212196.

9. Rubenstein, A.B., et al., Single-cell transcriptional profiles in human skeletal muscle. Scientific Reports, 2020. 10(1): p. 229.

10. Muto, Y., et al., Single cell transcriptional and chromatin accessibility profiling redefine cellular heterogeneity in the adult human kidney. Nature Communications, 2021. 12(1): p. 2190.

11. Wilson, P.C., et al., Multimodal single cell sequencing implicates chromatin accessibility and genetic background in diabetic kidney disease progression. Nature Communications, 2022. 13(1): p. 5253.

12. Lengyel, E., et al., A molecular atlas of the human postmenopausal fallopian tube and ovary from single-cell RNA and ATAC sequencing. Cell Rep, 2022. 41(12): p. 111838.

13. Genomics, x., PBMC from a Healthy Donor - Granulocytes Removed Through Cell Sorting (10k) - Single Cell Multiome ATAC + Gene Expression Dataset, x. Genomics, Editor. 2020: 10x Genomics.

14. Hao, Y., et al., Integrated analysis of multimodal single-cell data. Cell, 2021. 184(13): p. 3573–3587.e29.

