## Supplementary Figures for "Single-Cell Data Integration and Cell Type Annotation through Contrastive Adversarial Open-set Domain Adaptation"

† Equal first Authorship
* Corresponding Author

Correspondence to:

**Supplementary Figure 1.** Heatmap of confusion matrices illustrating SAFAARI's cell type predictions in 10X data, using FACS as the source dataset in an open-set domain adaptation scenario. The colour scale represents the percentage of cells classified each cell type (Page 2)

**Supplementary Figure 2.** UMAP visualizations of SAFAARI's integration across technologies (10X and FACS) for 8 tissues from Tabula Muris (Page 3).

**Supplementary Figure 3.** UMAP visualizations of SAFAARI's cell embeddings across 8 Tabula Muris tissues, demonstrating the integration of an unannotated query dataset into a reference dataset using pseudo-labels identified through label transfer (Page 4).

**Supplementary Figure 4.** UMAP visualization of learned cell embeddings and heatmap of the confusion matrix for cross-species open-set domain adaptation results on scRNA-seq muscle data. The analysis annotates human cell types using bovine (A) and mouse (B) as source domains, with monocytes treated as an unknown cell type (Page 5).

**Supplementary Figure 5.** Sankey plot and heatmap of the confusion matrix illustrating SAFAARI's label transfer from the normal kidney dataset to the DKD dataset, with T cells designated as the unknown class (Page 6).

**Supplementary Figure 6.** UMAP visualization comparing Harmony and Seurat integration of scRNA-seq and scATAC-seq batches of ovary dataset (Page 7).

**Supplementary Figure 1.**

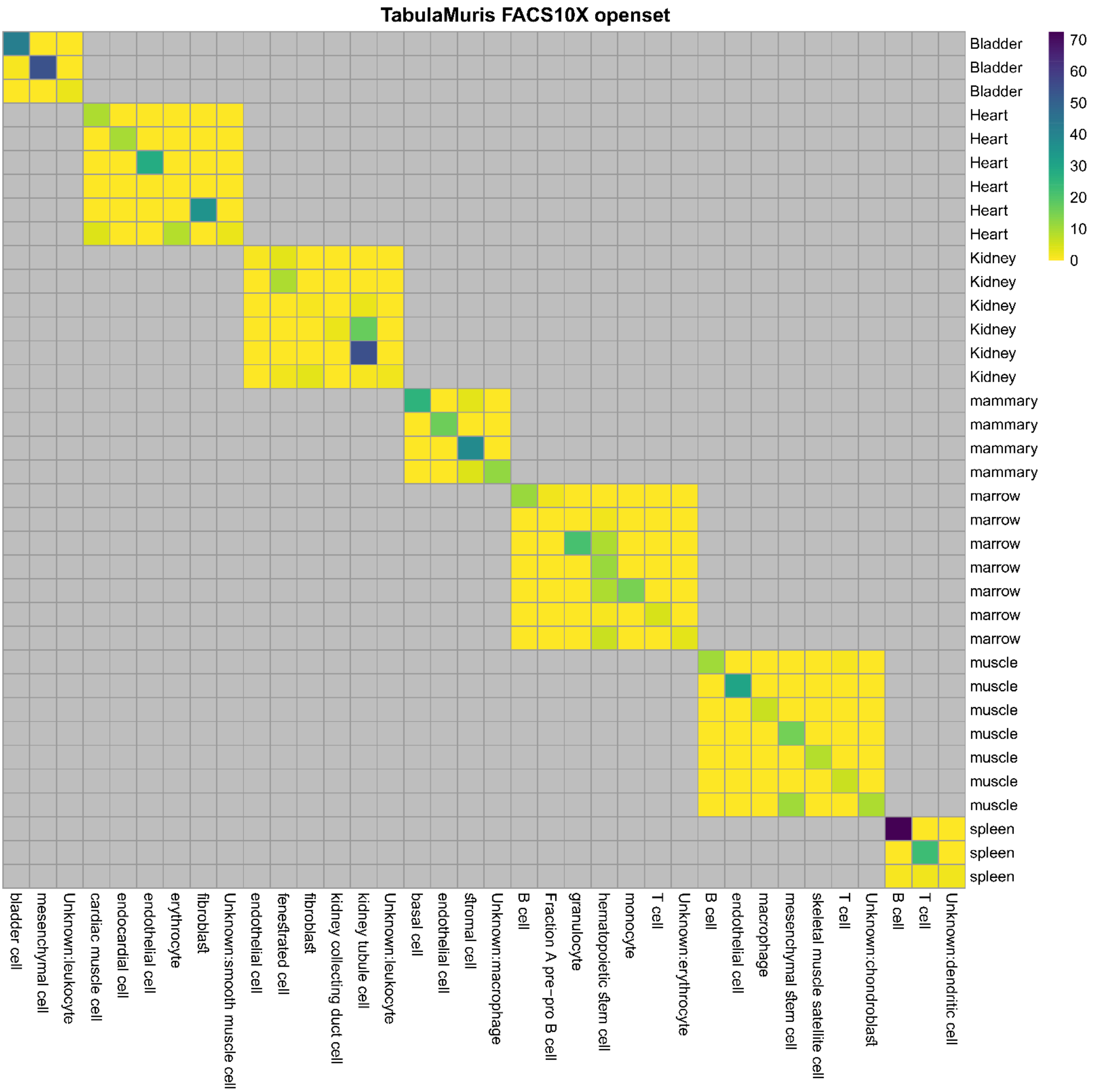

### Supplementary Figure 2.

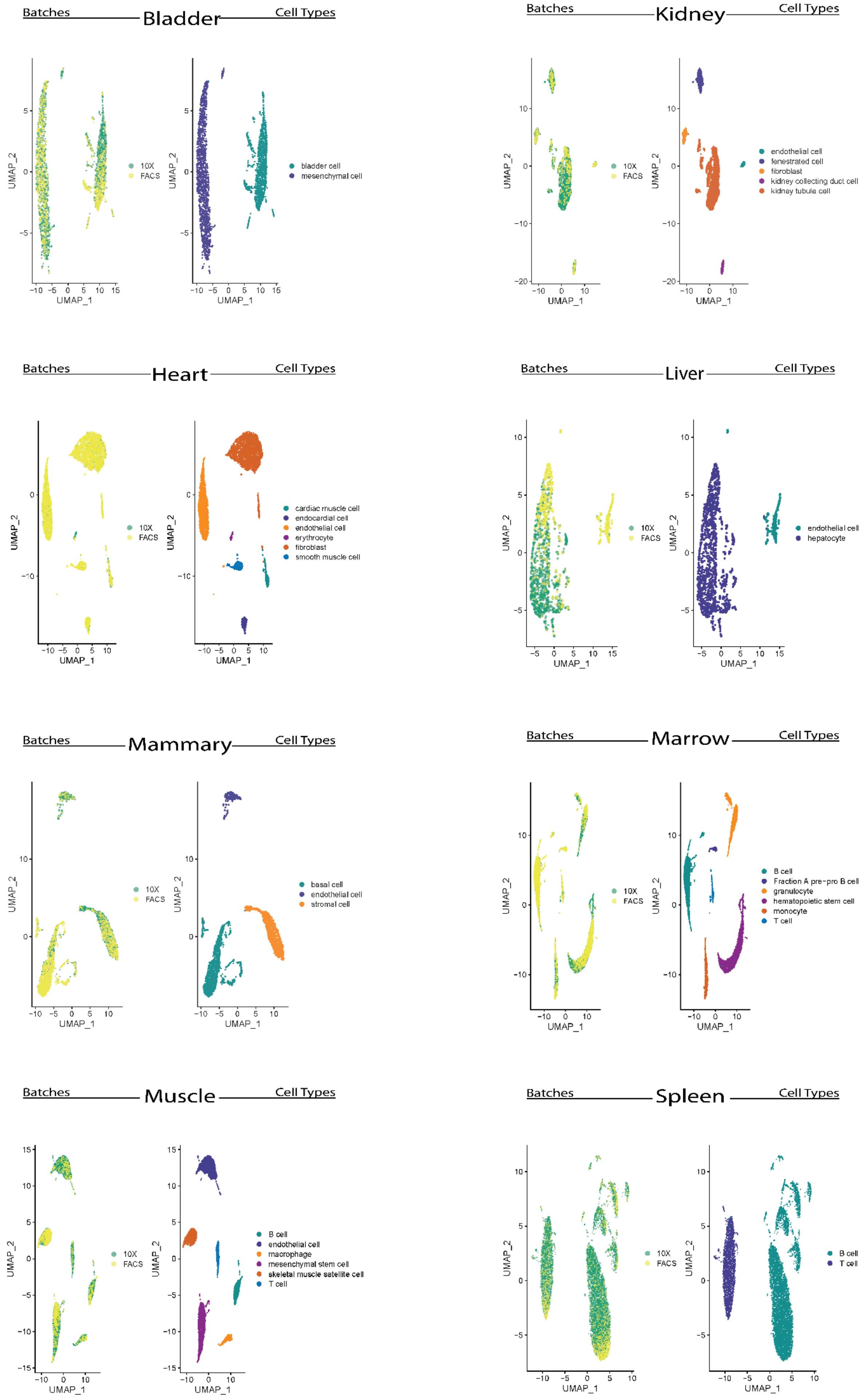

#### Supplementary Figure 3.

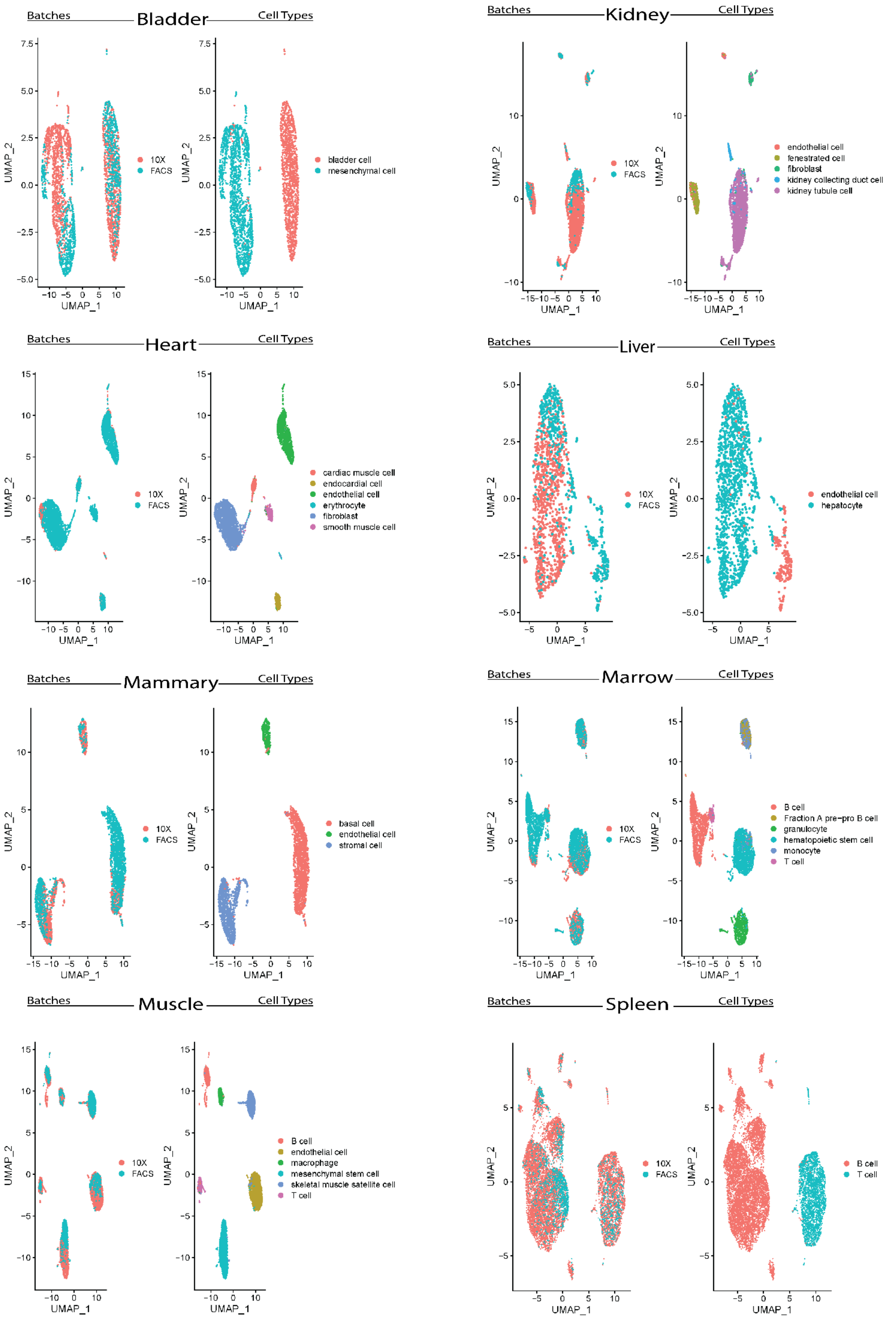

#### Supplementary Figure 4.

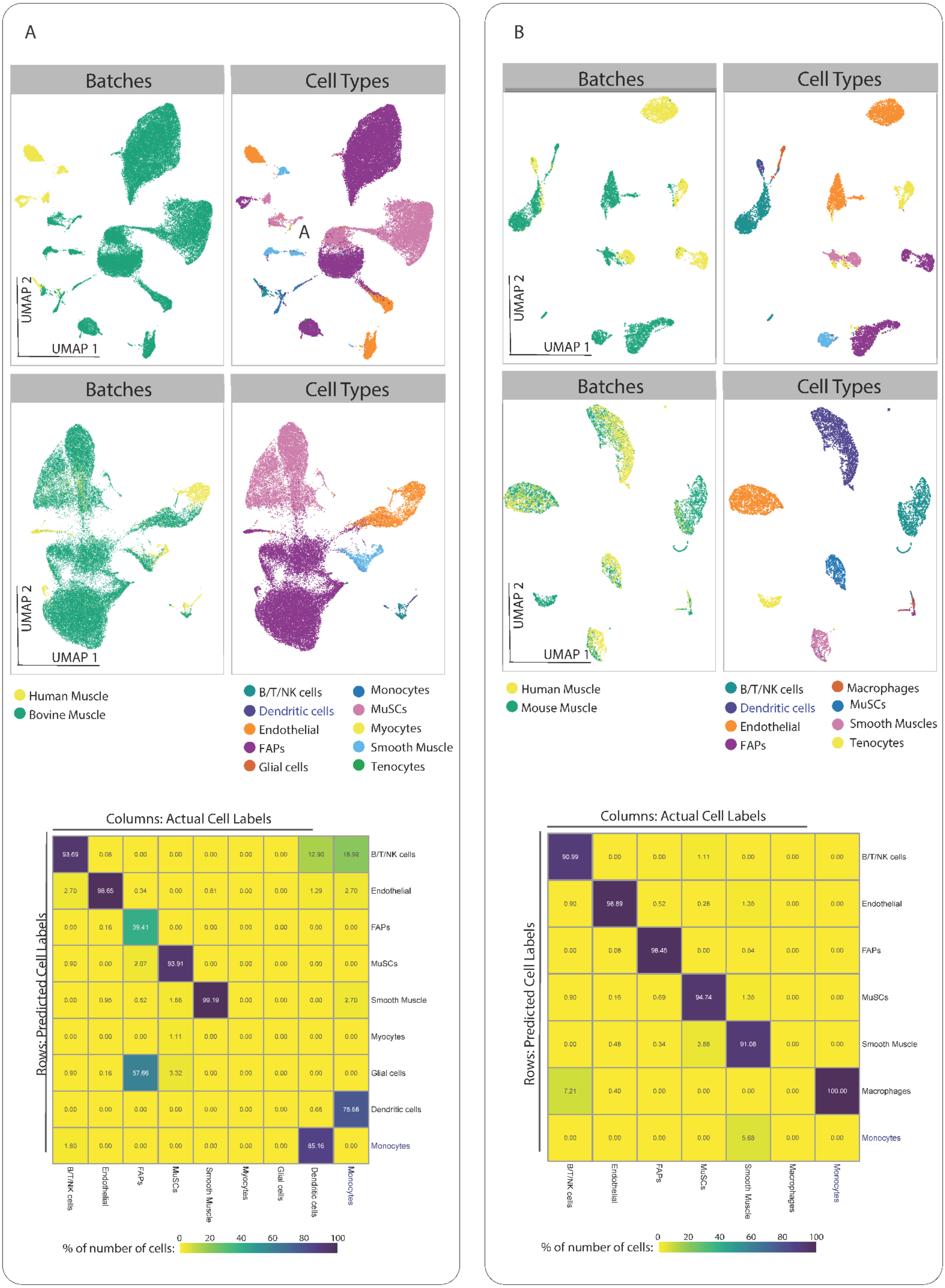

**Supplementary Figure 5.**

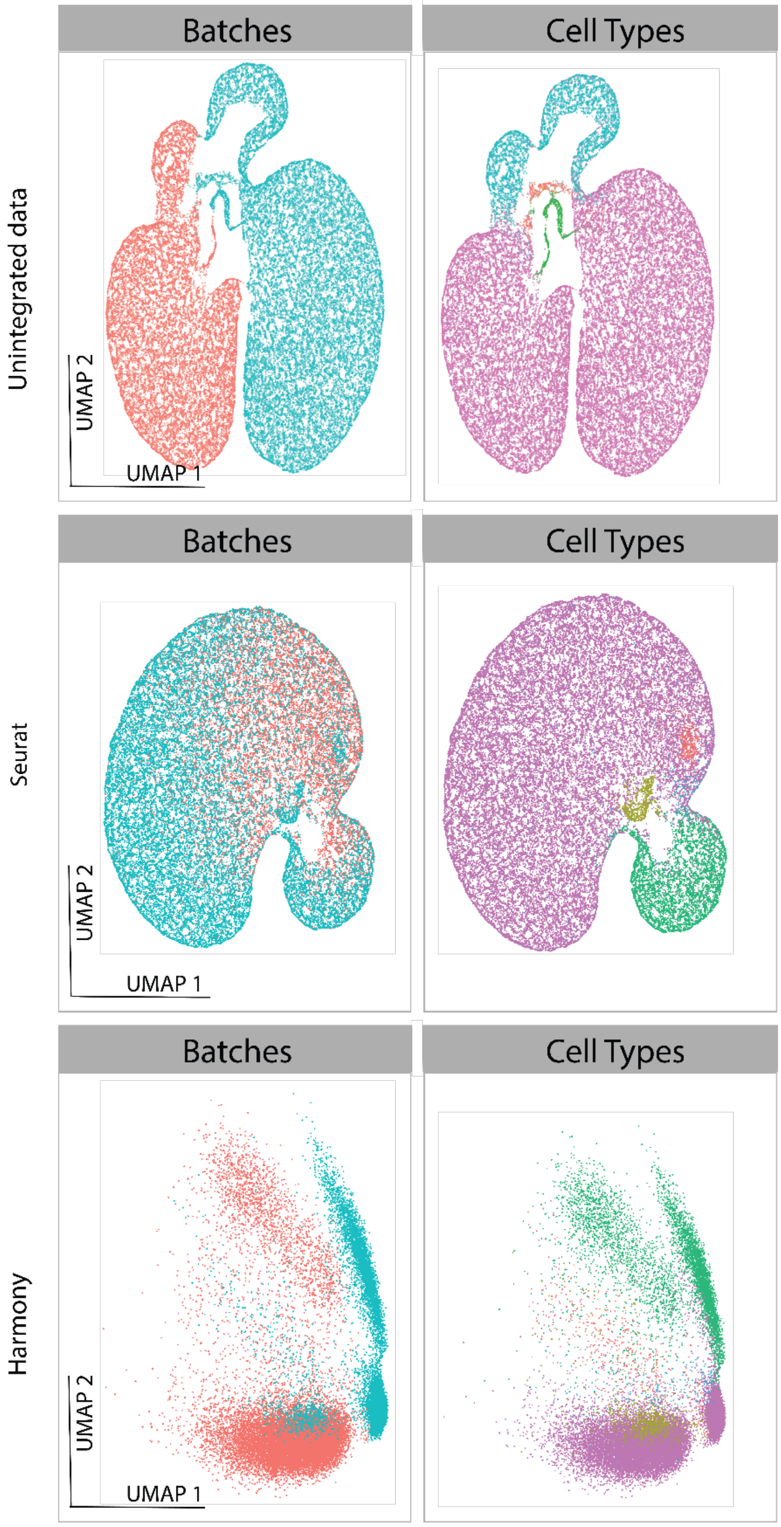

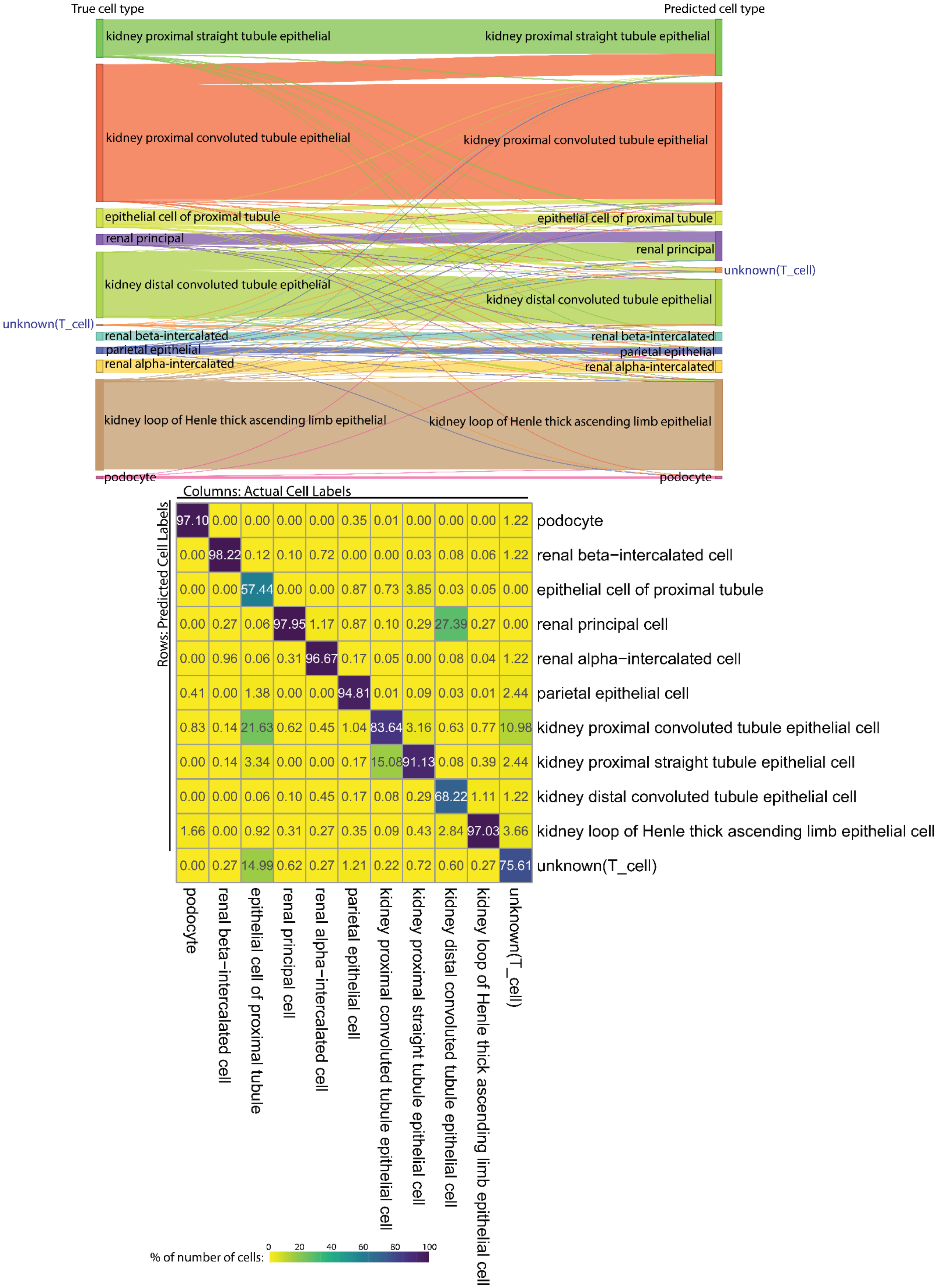


**Supplementary Figure 6.**

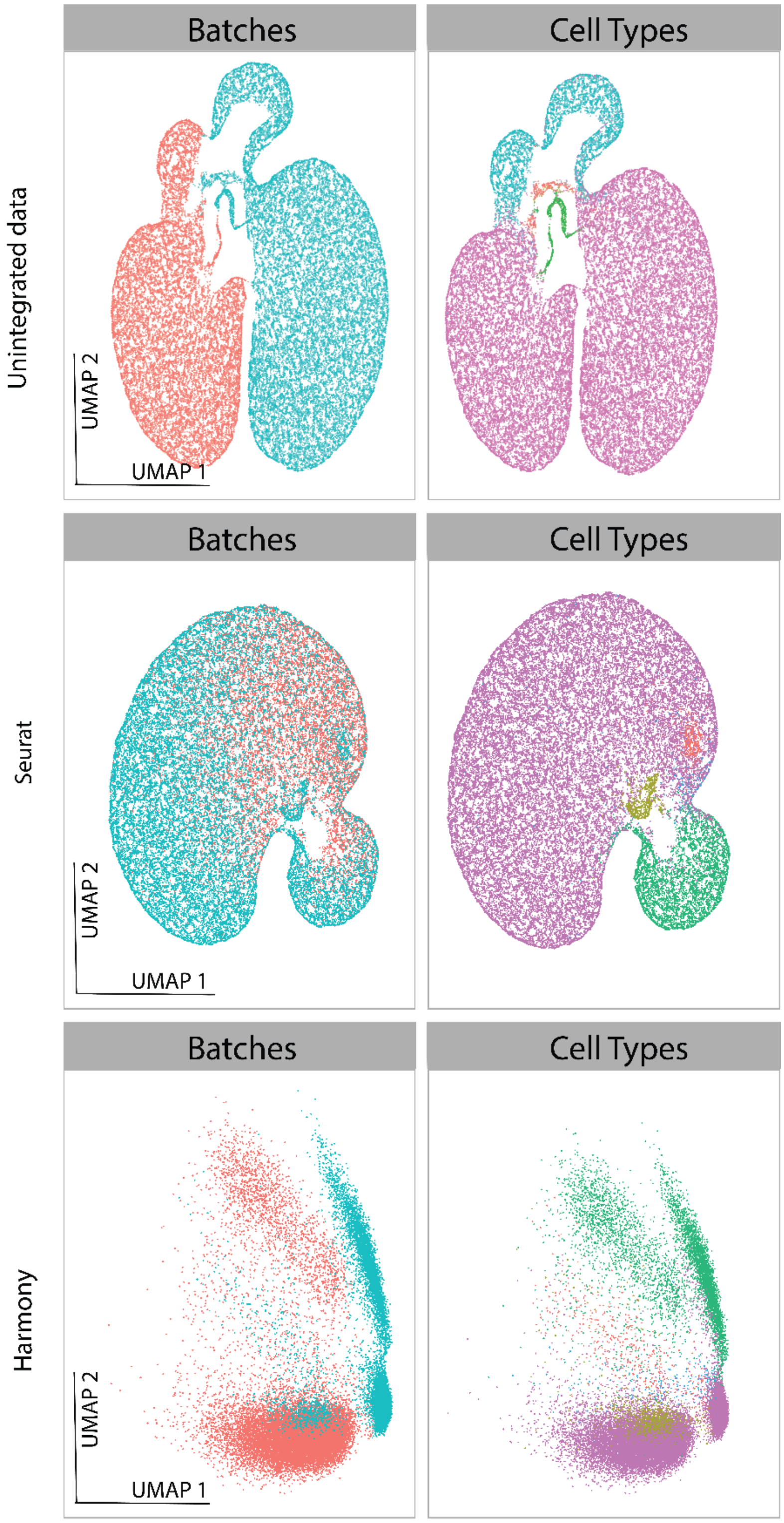
Batches
Cell Types
Unintegrated data
UMAP 2
UMAP 1
Batches
Cell Types
Seurat
UMAP 2
UMAP 1
Batches
Cell Types
Harmony
UMAP 2
UMAP 1
